# RNA gradients can guide condensates toward promoters: implications for enhancer-promoter contacts and condensate-promoter kissing

**DOI:** 10.1101/2024.11.04.621862

**Authors:** David Goh, Deepti Kannan, Pradeep Natarajan, Andriy Goychuk, Arup K. Chakraborty

## Abstract

We study how protein condensates respond to a site of active RNA transcription (i.e., a gene promoter) due to electrostatic protein-RNA interactions. Our results indicate that condensates can show directed motion towards the promoter, driven by gradients in the RNA concentration. Analytical theory, consistent with simulations, predicts that the droplet velocity has a non-monotonic dependence on the distance to the promoter. We explore the consequences of this gradient-sensing mechanism for enhancer-promoter (E-P) communication using polymer simulations of the intervening chromatin chain. Directed motion of enhancer-bound condensates can, together with loop extrusion by cohesin, collaboratively increase the enhancer-promoter contact probability. Finally, we investigate under which conditions condensates can exhibit oscillations in their morphology and in the distance to the promoter. Oscillatory dynamics are caused by a delayed response of transcription to condensate-promoter contact and negative feedback from the accumulation of RNA at the promoter, which results in charge repulsion.

## I. INTRODUCTION

Biomolecular condensates compartmentalize different biochemical reactions by enriching the local concentrations of specific reactants and enzymes^1,2^. Condensates are believed to form through liquid-liquid phase separation, which is driven by a network of weak multivalent interactions among proteins^1–4^. The properties of condensates can be modulated by interactions between proteins and nucleic acids, such as DNA and RNA^5–12^. When molecular interactions are coupled with irreversible chemical reactions, biomolecular condensates can exhibit a variety of rich phenomena not present in thermal equilibrium^13^. Past theoretical studies have investigated how reactions can lead to nonequilibrium phenomena such as arrested coarsening^14–21^, droplet division and anti-coarsening^18,20–22^, self-propulsion^20,23,24^, and directed motion^25–27^.

Coupling phase separation with irreversible chemical reactions may be a generic feature of intracellular dynamics. For example, transcriptional condensates are composed of proteins required for RNA transcription, including RNA Polymerase II, transcription factors (TFs), and coactivators^5,28–33^. These proteins have positively charged regions which interact electrostatically with the freshly transcribed negatively charged RNA^12^. These interactions were shown to give rise to a reentrant phase transition, where at low RNA concentrations, attractive interactions recruit proteins into the condensate, whereas at high RNA levels, repulsive interactions expel proteins from the condensate^10–12^. As a result, an excess of transcribed RNA can dissolve transcriptional condensates, providing a negative feedback loop for transcription^12^. When transcription is spatially localized to active genes, computer simulations further show that condensates can move toward these genes and adopt nonequilibrium steady state morphologies, such as vacuoles and aspherical shapes^27^. These simulation results are corroborated by experimental studies that have observed nuclear condensates showing directed motion^34–37^ and adopting aspherical morphologies^38^.

Past theoretical studies have characterized the motion of condensates due to externally imposed ^25,26^ or self-generated concentration gradients^20,24^. In both cases, droplet motion is driven by a chemical potential gradient across the droplet, which creates a net flux of proteins^24^. Motivated by these theoretical results, and experimental observations of droplet motion *in vitro*^39,40^, we asked if this *gradient-sensing* mechanism could explain the localization of transcriptional condensates towards active gene promoters, which generate RNA concentration gradients via transcription. Building on the self-consistent sharp interface theory developed in Ref.^24^, we analytically calculate the velocity of transcriptional condensates as they flow towards gene promoters and demonstrate its agreement with simulations. We then explore the consequences of this directed motion for gene regulation, focusing on the interactions between enhancers, condensates, and promoters.

Transcriptional condensates form at chromatin loci enriched with TF-DNA binding sites, such as superenhancers^5,28,29,41^, and are believed to remain bound to chromatin^30^. Enhancers regulate gene expression by interacting with their cognate promoters ^42^, but the mechanisms that bring enhancers and promoters into spatial proximity remain unclear^43^. Cohesin can increase the contact frequency between enhancers and promoters by extruding loops^44^, yet depleting cohesin does not fully abrogate all enhancer-promoter contacts^45–52^. Transcriptional condensates are speculated to act as a bridge between the enhancer and the promoter, a mechanism which is known as the “action-at-a-distance” model^43,53,54^. As a result, the condensate enriches the concentration of transcriptional proteins around the promoter, which may drive higher rates of transcription^28,32,55^. However, the role of condensates in bringing the enhancer and promoter into proximity, if any, remains unexplored^56^.

Here, we propose that condensate motion in response to an RNA concentration gradient results in a non-reciprocal interaction which attracts enhancer-bound condensates toward promoters, but not vice versa. Brownian dynamics simulations of a polymer model of chromatin show that this non-reciprocal interaction increases the enhancer-promoter contact probability. We hypothesize, supported by simulations, that gradient-sensing can cooperate with loop extrusion to robustly give rise to enhancer-promoter contacts over different length scales^57^.

We next shift our focus to more dynamic models of condensate-promoter communication. In particular, live-cell imaging studies have observed transient and dynamic kissing events between condensates and gene promoters reminiscent of oscillations^30,33^. Inspired by these observations, we wondered under what conditions our model could admit oscillations in condensate-promoter proximity. In the cell, the transcription machinery enriched within transcriptional condensates first assembles into a preinitiation complex. RNA polymerase II then stalls near the promoter during promoter-proximal pausing, before finally proceeding to the productive elongation of RNA^58–60^. These processes can give rise to an effective time delay between condensate-promoter contact and transcription. We demonstrate that coupling such delays with the negative feedback from RNA accumulation^10–12^ can produce oscillatory dynamics. Finally, we discuss the implications of our results for dynamic models of enhancer-promoter communication.

## II. MODEL

Based on prior studies^12,27,61^, we consider a phase field model where a conserved species of transcriptional proteins with concentration *c*(***x***, *t*) regulates the production of a non-conserved species of nascent RNA, with concentration *m*(***x***, *t*). We assume that minimization of the following free energy of protein-protein interactions results in their spontaneous phase separation at equilibrium:

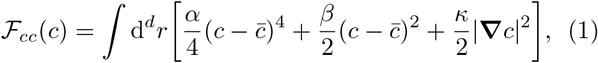

where 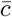 is the critical point. The parameter *β <* 0 sets the difference between the concentrations of the protein-rich and the protein-poor binodal points, 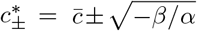. The surface tension, which acts to minimize concentration gradients, is set by the parameter *κ >* 0.

Electrostatic interactions between positively charged proteins and negatively charged RNA lead to a re-entrant phase transition driven by a balance of charges, which is known as complex coacervation^10–12^. This phenomenon can be captured using a phenomenological free energy of the form

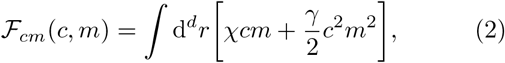

where attractive interactions (*χ <* 0) dominate at low RNA concentrations, and repulsive interactions (*γ >* 0) dominate at high RNA concentrations. Although symmetry arguments do not preclude terms of the form *mc*^2^ and *cm*^2^ in the above Landau-Ginzburg expansion, it was previously shown that these terms are not necessary for the system to show a reentrant phase transition^12^. In Sections III and IV we first focus on the linear order term (i.e., we make the approximation *γ* ≈ 0), and later explore the effects of repulsive RNA-protein interactions in Sec. V. Altogether, the thermodynamics of the protein-RNA system is characterized by the free energy functional,

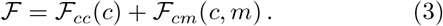

Since protein turnover is slow compared to nascent RNA transcription, we assume the amount of protein to be conserved. Protein mass is redistributed by currents driven by gradients in the net free energy,

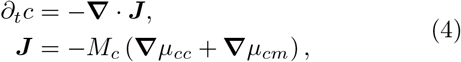

where we have separated the contributions to the current due to protein-protein interactions alone, *µ*_*cc*_ = *δℱ*_*cc*_*/δc*, and due to protein-RNA interactions, *µ*_*cm*_ = *δℱ*_*cm*_*/δc*. The protein mobility *M*_*c*_ is assumed to be constant for consistency with prior studies^12,27,61^.

RNA is actively produced by proteins via the consumption of nucleoside triphosphates (NTPs) in regions with high concentrations of the transcription machinery, with the rate *k*_*p*_*c >* 0 spatially localized around a point on the gene locus. The dynamics are thus governed by a reaction-diffusion equation,

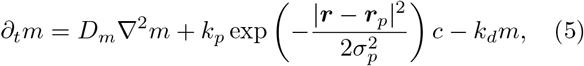

where *D*_*m*_ is the diffusion coefficient of nascent RNA, and *k*_*d*_ is the rate at which nascent RNA is modified into a form that ablates its interactions with transcriptional proteins. Since the DNA from which the RNA is transcribed has a Gaussian end-to-end distribution, the variance 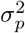 of the spatial production rate is a proxy for the length of the gene. Alternatively, the Gaussian localization function can be interpreted as the density of promoters in a cluster of genes^27^. We simulate the coupled protein and RNA dynamics with the promoter located at the origin (***r***_*p*_ = **0**). We employ finite volume simulations using a circular mesh with no-flux boundary conditions for both species. We set the length scale to be the diffusion length of RNA, 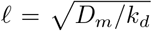, and the time scale to be the inverse of the RNA degradation rate, 1*/k*_*d*_ (Supplementary Information, Sec. SV).

## III. RNA GRADIENT SENSING CAUSES DIRECTED MOTION OF CONDENSATES TOWARDS GENE PROMOTERS

We investigate how a condensate responds to RNA concentration gradients generated at an active promoter (Fig. 1A) using simulations and theoretical analysis of the dynamics in Eqs. (4) and (5). In the following, we consider a regime where the RNA-protein interactions are attractive. That is, we only include the linear order term in Eq. (2), *χcm*, in order to ease comparisons to analytical theory. In our phase field simulations, we initialize a circular droplet of radius *R*(*t* = 0) = 1 with initial dense phase concentration *c*_+_(*t* = 0) = 5.5 at a fixed initial distance *r*(*t* = 0) = 10 between the condensate center-of-mass and the promoter, and track the protein and RNA dynamics via a finite-volume method simulation of Eqs. (4) and (5). We set the initial RNA concentration to be *m*(***r***, *t* = 0) = 0.

**FIG. 1.**
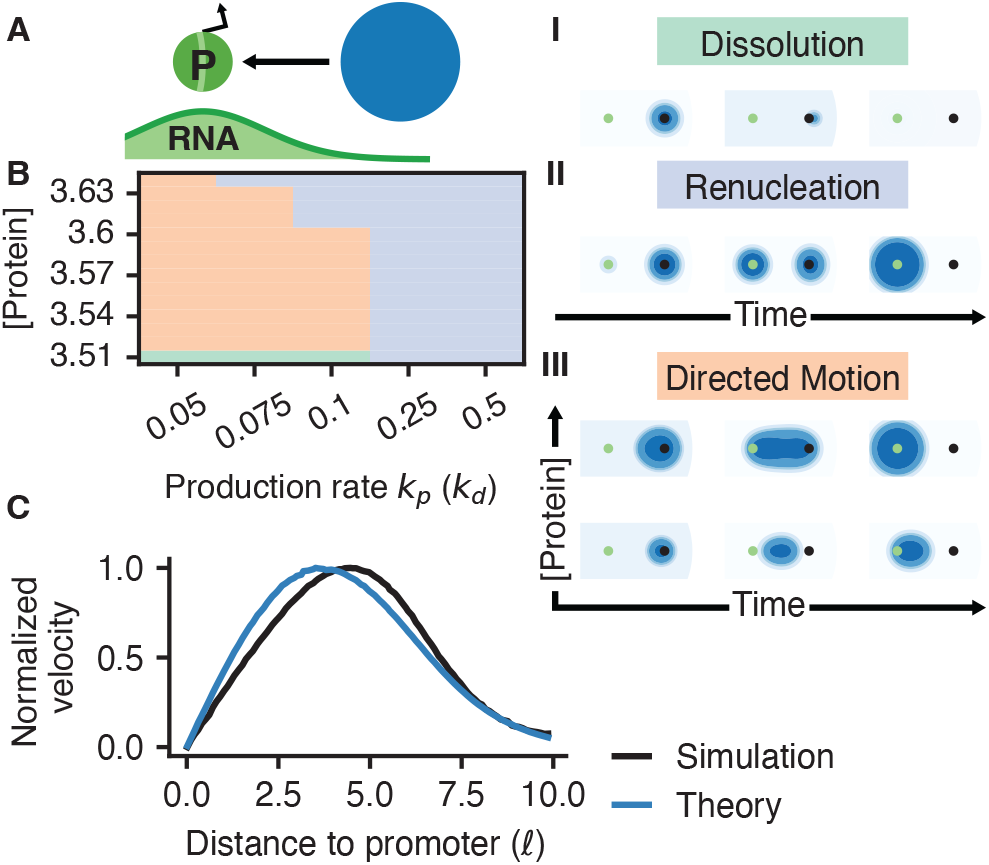
Condensates can sense RNA gradients and move towards gene promoters. (A) Transcription at the gene promoter produces RNA, which diffuses and degrades, resulting in an RNA concentration gradient. (B) Condensates can dissolve (I), dissolve and renucleate at the promoter (II), or exhibit directed motion towards the promoter (III) depending on the RNA production rate *k*_*p*_ (in units of the degradation rate *k*_*d*_) and the total protein concentration, which we control by varying the initial concentration of proteins outside the droplet *c*_(*t* = 0). Simulation parameters: *α* = 1, *β* = − 0.25, *κ* = − 0.1, *χ* = −0.1, *γ* = 0, *M*_*c*_ = 1, *D*_*m*_ = 1, *σ*_*p*_ = 2.5, *k*_*d*_ = 1, *c*_+_(*t* = 0) = 5.5, *m*(***r***, *t* = 0) = 0, *r*(*t* = 0) = 10, and *R*(*t* = 0) = 1. (C) Droplet velocity, normalized by the maximum value, is non-monotonic in the distance between the droplet center of mass and the promoter (reported in units of the RNA diffusion length, 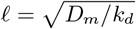. The theory underpredicts the maximum of the velocity by 29.3%. Theoretical predictions show that directed motion is driven by RNA gradients [Eq. (9)]. Simulation parameters specific to panel (C): *k*_*p*_ = 0.08 and *c*_(*t* = 0) = 3.53.

We find that the dynamics of the droplet are most influenced by the RNA production rate and the total amount of protein in the system, which we control by varying the initial concentration of proteins outside the droplet (Fig. 1B). For average protein concentrations close to the protein-poor binodal point 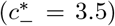, the nucleated droplet dissolves because a homogeneous light phase admits a lower free energy than a two-phase system (Fig. 1B I). At high RNA production rates, the protein-RNA attraction at the promoter is strong enough to dissolve the current condensate and nucleate another (Fig. 1B II). At lower RNA production rates, if the protein concentration is not too low, the droplet instead flows towards the RNA source (Fig. 1B III). The droplet elongates as it flows, as has been observed in experiments on the directed motion of nuclear speckles^37^. Droplet elongation is enhanced at higher protein concentrations (Fig. 1B III).

To further investigate what drives droplet flow, we quantify the flow velocity analytically with the method outlined in Ref.^24^. First, we assume the droplet flows with a constant velocity (i.e., steady motion), and thus seek a traveling-wave solution *c*(***x − v****t*) ≡ *c*(***z***) to the continuity equation,

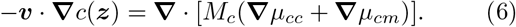

To analyze Eq. (6) analytically, we take the sharp interface limit, meaning the interface width 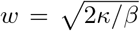 is smaller than all other length scales relevant to the dynamics. Mathematically, we simultaneously take the limits *α* → ∞ and *β* → ∞, which maintains finite dense and light phase concentrations, 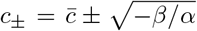 while implying an infinite effective surface tension and therefore a round droplet shape^24^. We thus consider a piecewise constant protein concentration profile *c*(***z***) where the concentration within the droplet is assumed to be equivalent to the protein-rich binodal, 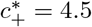, and the concentration outside the droplet is set to the protein-poor binodal, 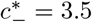.

Having imposed a droplet concentration profile, we temporarily assume knowledge of the droplet velocity ***v*** and derive a constraint on *µ*_*cc*_(***z***| ***v***) that adiabatically maintains a piecewise constant protein concentration field. In Eq. (6), the effective force due to protein-RNA interactions, −∇*µ*_*cm*_, breaks energy conservation because RNA is continuously produced and degraded. In contrast, the current of proteins ***j*** = *c*(***z***)***v*** in the presence of the purely thermodynamic force −∇*µ*_*cc*_ should not dissipate power^24^,

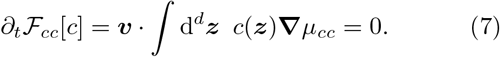

Because *µ*_*cc*_ far from the droplet is uniform, following Ref.^24^, one can show that Eq. (7) reduces to the following thermodynamic consistency condition involving an integral over the domain 𝒟 of the droplet, |𝓏| ≤ *R*,

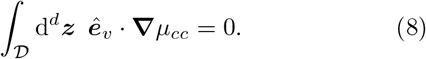

The condition in Eq. (8) implies that the effective force due to protein-protein interactions alone must be symmetric about the droplet center in the direction of flow ***ê*** _*v*_.

With the above simplifications, one can first solve Eq. (6) for the unknown function *µ*_*cc*_(***z*** | ***v***) and then use Eq. (8) to determine the droplet velocity ***v***^24^. For a constant mobility of the proteins, we find that in *d* = 2 or *d* = 3 dimensions (Supplementary Information, Sec. SIII):

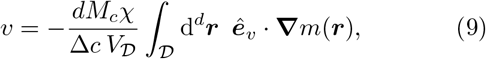

where *V*_*D*_ ∼ *R*^*d*^ is the droplet volume. Equation (9) demonstrates that the directed motion of the droplet is driven by concentration gradients in RNA. The velocity is proportional to the strength of the RNA-protein attraction *χ* and the protein mobility *M*_*c*_, but inversely proportional to the concentration difference between the dense and light phase, 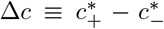. Note that the droplet velocity is enhanced by a factor of the number of dimensions, *d*, compared to calculations reported in prior literature where the droplet is treated akin to a solid so that all material points within the droplet move at the same velocity^25,27^. In essence, transport is driven both by a roughly 1D flux of proteins in the dense phase, and, in addition, amplified by *d*− 1 dimensions in which droplet material is brought from the trailing edge across the light phase and into the leading edge of the droplet. Thus, the leading edge is continuously being renucleated as the droplet moves, which explains why the droplet velocity is inversely proportional to the strength of phase separation, Δ*c*.

To test our theoretical calculations, we assume that the RNA concentration field reaches steady state quickly compared to the dynamics of the droplet and, moreover, that this steady-state profile is unaffected by the droplet. We thus obtain the fixed RNA profile from the steady state solution of Eq. (5) and substitute the RNA concentration gradient in Eq. (9). Equation (9) also assumes a fixed droplet radius; however, in simulations, the droplet radius grows with time due to coarsening and attractive interactions with the RNA, which promote incorporation of the light phase protein into the dense phase^25^. In the absence of an analytical expression for the droplet size and shape as a function of time, we determine its effective radius from simulations by calculating the angular average of the distance between each point along the contour of the droplet and the droplet center of mass (Supplementary Information).

In Fig. 1C, we compare the velocity, normalized by the maximum value, of simulated droplets as a function of distance from the RNA source, *r*, to the theoretical prediction of Eq. (9). When the droplet centroid and the RNA source colocalize, the RNA concentration profile is symmetric about the droplet center, causing motion to arrest. In the range *r* ≤ *R*, the gradient in the RNA concentration profile progressively increases, until it reaches a maximum at *r* = *R* = 4, where the leading edge of the droplet coincides with the maximum of the RNA profile. In our simulations, the droplets elongate as they flow; as a result, the distance at which the leading edge coincides with the RNA source is the major axis of the ovular droplet, which is larger than the angular average of the droplet radius (Supplementary Information, Fig. S4) used in the theoretical calculation. Thus, in simulations, the velocity peaks at a distance slightly larger than the droplet angular average radius. In simulations of perfectly round droplets, the peak of the simulation and theory curves align (Supplementary Information, Fig. S7).

While the shape of the velocity versus distance curve is similar in simulations and theory, the theory underpredicts the maximum of the velocity by 29.3%. One reason for the discrepancy is that the simulations in Fig. 1C include a time-varying RNA gradient which responds to droplet motion. Simulations of droplet motion in response to a fixed RNA gradient more closely match the theoretical prediction (Supplementary Information, Fig. S2). In addition, our theoretical calculations assume Δ*c* = 4.5− 3.5 = 1 is determined by the binodal points, whereas simulations show that the difference in concentration between the dense and light phase is always less than one and increases as the droplet flows towards the promoter. Accounting for both of these factors explains most of the discrepancy between simulations and theory (Supplementary Information, Fig. S2).

## IV. DIRECTED MOTION OF ENHANCER-BOUND CONDENSATES CAN INCREASE ENHANCER-PROMOTER CONTACTS

Overall, our theory and simulations show that RNA concentration gradients can lead to the long-ranged attraction of protein droplets towards an actively transcribing gene promoter. We next asked whether the directed motion of condensates, in collaboration with loop extrusion by cohesin, would influence the proximity of the enhancer and the promoter. We consider the dynamics of the intervening chromatin polymer, which we model as a Rouse chain^62^. Despite neglecting many details of chromatin structure, Rouse models have generated many insights into polymer dynamics and have been shown to recapitulate dynamical data on chromosomes^57,63,64^.

The 3D distances between two loci connected by a Rouse chain of length *L*_*C*_ follow a Maxwell distribution *P* (*r*) = 4*πr*^2^(3*/*(4*L*_*c*_𝓁_*p*_*π*))^3*/*2^ exp(−3*r*^2^*/*(4*L*_*c*_𝓁_*p*_)) where 𝓁_*p*_ is the persistence length of chromatin. If there are non-overlapping, non-interacting loops in between the two loci, the intervening distance distribution has a reduced effective contour length *L*_*C*_^65^. Live-cell imaging experiments estimate that loop extrusion with boundaries reduces the effective genomic separation between two CTCF loci by 61%^57^. Thus, rather than simulating the actual loop extrusion process, which requires introducing several parameters such as the extrusion speed and residence time of cohesin on DNA^66^, in this work we simply assume a reduced contour length to model the effect of cohesin on enhancer-promoter distances.

Active transcription at a gene promoter results in an RNA concentration gradient, which attracts an enhancer-bound condensate towards the promoter, but not vice versa. If we assume the condensate is strongly tethered to the enhancer locus, the enhancer should move at the same velocity as the condensate. In 3D, the velocity of a spherical condensate of radius *R* at a distance *r* away from a point source of RNA has a closed analytical expression (Supplementary Information, Sec. SIII),

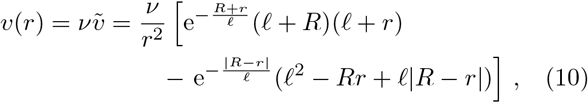

where 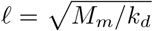 is the diffusion length of RNA and *ν* sets the velocity scale,

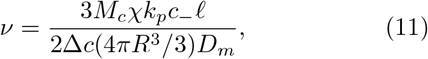

such that 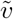 is dimensionless. As seen in Fig. 2B, the dimensionless velocity is sharply peaked at the droplet radius *r* = *R*. It falls off as *r*^−2^, mirroring the gradient in the RNA profile. Here, we use a droplet radius of *R* = 250 nm based on measurements of transcriptional condensates *in vivo*^30,33^ and estimate the RNA diffusion length to be 𝓁 = 4.24 µm (Supplementary Information, Sec. SVIII). The magnitude of the velocity, *ν*, additionally depends on gene-specific parameters, such as the production rate *k*_*p*_. We thus treat *ν* as a parameter that we vary in our Brownian dynamics simulations.

**FIG. 2.**
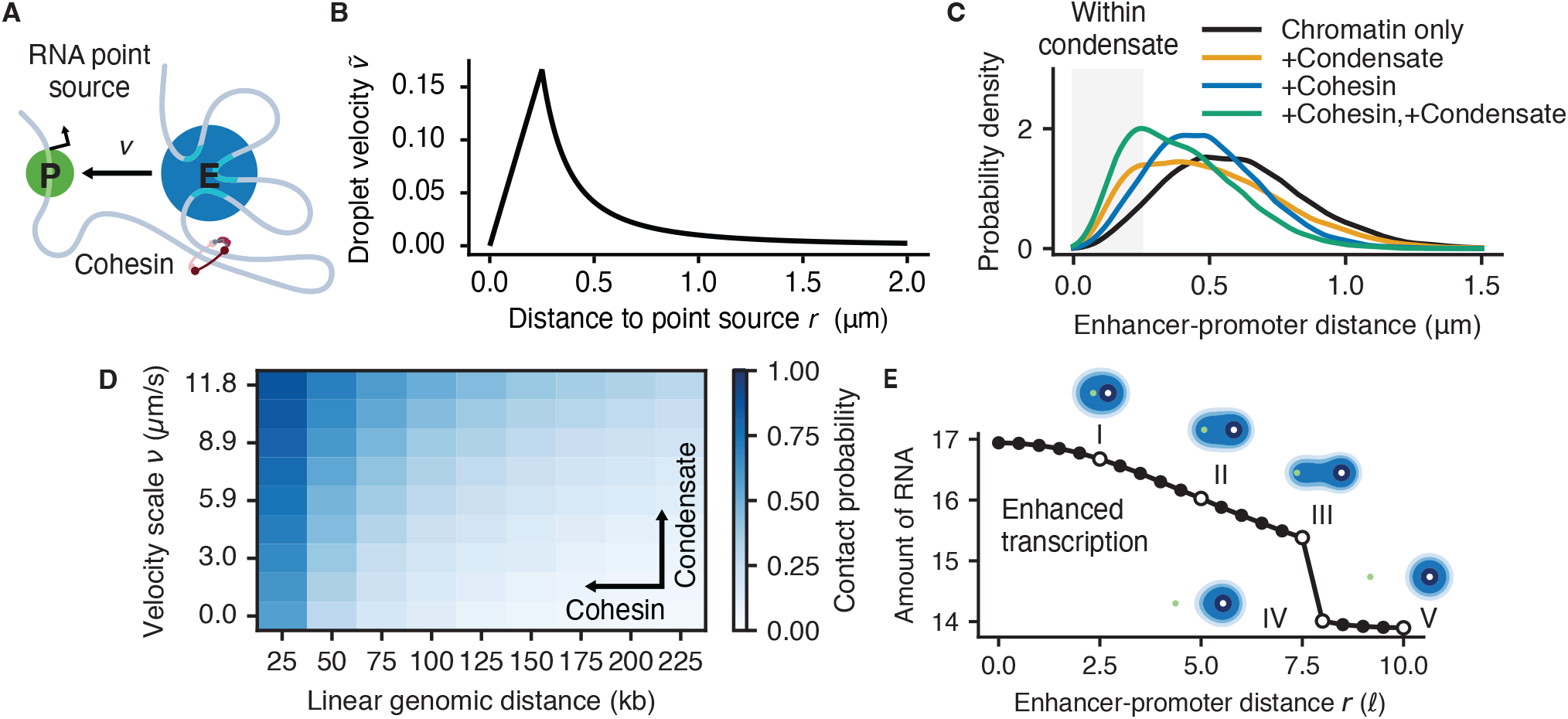
Directed motion of enhancer-bound condensates can contribute to enhancer-promoter proximity, and hence, enhanced transcription at the promoter. (A) We calculate the distribution of enhancer-promoter separations using a polymer model that takes into account loop extrusion by cohesin and the directed motion of enhancer-bound condensate towards the promoter. (B) Dimensionless droplet velocity in response to an RNA point source in 3D [Eq. (10)], which resembles an effective attractive force moving the enhancer toward the promoter [Eq. (12)]. Here, we use a droplet radius of *R* = 250 nm based on measurements of transcriptional condensates *in vivo*^30,33^ and estimate the RNA diffusion length to be 𝓁 = 4.24 µm (Supplementary Information, Sec. SVIII). (C) Distribution of separations between an enhancer and promoter that are 150 kilobases apart. The presence of an enhancer-bound condensate and transcription at the promoter shifts the distribution to smaller separations (in this example, the velocity prefactor is *ν* = 10 µm s^−1^). Cohesin also brings enhancers closer to promoters by reducing the effective contour length of the intervening polymer (in this example, by a factor of 2/3). Both of these effects collaborate to bring the enhancer within a condensate radius of the promoter. (D) The enhancer-promoter contact probability as a function of the contour length of chromatin connecting the two loci and the magnitude of the condensate velocity towards the promoter. We define the contact probability as the probability that the enhancer-promoter distance is within the radius of the condensate, assumed to be 250 nm. (E) For a fixed enhancer-promoter separation, our simulations show that RNA transcription is enhanced at the promoter when the condensate overlaps with the gene promoter.

Incorporating a non-reciprocal force into the Rouse chain dynamics will lead to probability currents and a steady state that cannot be described by a Boltzmann weight. To calculate this non-equilibrium steady state distribution, we perform Brownian dynamics simulations of a discretized Rouse chain, where each monomer represents a Kuhn segment of length *b* = 2𝓁_*p*_. We choose a Kuhn length of 35.36 nm (441.42 bp) based on predictions from a “zig-zag” polymer model of nucleosome-bound DNA in mouse embryonic stem cells^67^. In the absence of measurements detailing how enhancer DNA is organized within the condensate, we coarse-grain the enhancer locus as a single monomer, indicated by the index *e*. Under these assumptions, the dynamics of the monomers *n* ∈ [1, *N* ] connected by Hookean springs *k* is given by

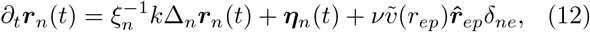

where Δ_*n*_ is the discrete Laplacian. The enhancer persistently moves in the direction of the promoter, 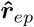, with a velocity which depends on the distance *r*_*ep*_ between the enhancer and the promoter. The random excitations ***η***_*n*_(*t*) have zero mean and in general have a covariance that depends on the friction *ξ*_*n*_ and activity *A*_*n*_ of each locus^68^,

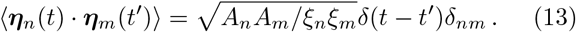

We assume that all monomers except for the enhancer have identical activity and friction with average diffusion coefficient *D*_*c*_ = *A*_0_*/ξ*_0_; however, we expect that the presence of a large condensate of radius *R* = 250 nm at the enhancer would increase *ξ*_*e*_ by a factor of 2*R/b* ≈ 14. Different choices for the friction and activity of the enhancer do not qualitatively change our results (Supplementary Information, Fig. S16). In our simulations, we set *b* = 1 and *D*_*c*_ = 1 for simplicity, but convert to experimental length and time scales using the Kuhn length *b* = 35.6 nm and the apparent diffusivity of chromatin *D*_app_ = 0.01 µm^2^*/*s^1*/*2^ (Supplementary Information, Sec. SVIII).

In the following, we assume that the RNA concentration gradient and enhancer-bound condensate are stable on the timescales for the polymer dynamics to reach steady state. This assumption is reasonable, since the average mammalian gene takes 10-20 minutes to transcribe^69^, and condensates localized to super-enhancers have been reported to remain stable for a similar timescale^30^. In contrast, we estimate the Rouse time for a 100 kbp chain to be on the order of 10 seconds. Nonetheless, condensate-promoter contact could alter transcription rates at the promoter and could also potentially dissolve or repel the condensate. For the purposes of this work, we neglect such feedback loops and calculate the steady-state enhancer-promoter distance distribution from an ensemble of 2000 snapshots across 200 independent simulations.

Cohesin-mediated loop extrusion effectively reduces the enhancer-promoter (E-P) tether length, resulting in a decreased mean enhancer-promoter distance (Fig. 2C, blue line). The velocity of the condensate is largest in a finite range near the condensate’s radius (Fig. 2B). Hence, the directed motion of the condensate shifts the probability density from large enhancer-promoter distances to distances that are within the condensate’s radius (Fig. 2C, yellow line). Our simulations show that cohesin and the condensate can collaboratively increase the probability of enhancer-promoter contacts (Fig. 2C, green line). We find that the contact probability decreases with increasing linear genomic distance, which implies that it increases in the presence of cohesin (Fig. 2D). In addition, it increases with the condensate velocity *ν* (Fig. 2D), which is proportional to the transcription rate at the promoter *k*_*p*_ [Eq. (11)]. Due to the finite range in which the condensate velocity is maximized, the contact probability is less sensitive to changes in the condensate velocity *ν* for larger genomic distances (Supplementary Information, Fig. S15). In contrast, cohesin would on average compact any genomic enhancer-promoter distance by the same factor, which we assume to be 2/3 based on live-cell imaging studies^57^.

We next wondered what the consequence of increased contact probability is on transcription at the promoter locus. To investigate this question, we return to our phase field simulations where the transcription rate is proportional to the protein concentration. The condensate may be “tethered” to the enhancer locus because of protein-mediated interactions. To model this approximately, we introduce a term to Eq. (3) which represents an attractive Gaussian potential toward the enhancer at a fixed position ***r***_*e*_.

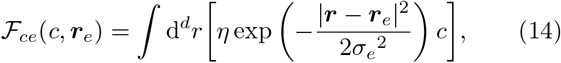

and initialize the droplet at the enhancer. Note that the use of a potential in the free energy is an approximation for a possibly non-equilibrium driving force for the condensate to be “tethered”. We find that the amount of RNA produced at the promoter increases sharply below a fixed enhancer-promoter distance (Fig. 2E). This long-ranged effect of an enhancer on transcription has been described in the literature as the “action-at-a-distance” model^43^. For a perfectly round droplet, the enhancer-promoter distance at which transcription is enhanced is simply the radius of the condensate. In our simulations, the droplet elongates, which increases the enhancer-promoter distance below which transcription is enhanced (Fig. 2E). The droplet elongates when the attraction of protein towards RNA at the promoter-proximal interface outcompetes the attraction of the droplet to the enhancer as well as surface tension.

According to the “action-at-a-distance” model, we expect the effective transcription rate to be linear in the contact probability, which increases with cohesin and the condensate. However, past experimental studies have shown how transcription can have a nonlinear dependence on enhancer-promoter contact probability^42^. In other words, once a threshold in contact probability is exceeded, further increases in contact probability would not have a significant effect on transcription. For a fixed condensate velocity *ν*, enhancer-promoter pairs at small genomic separations already have a contact probability that exceeds this threshold, independent of cohesin-mediated compaction (Fig. 2D). However, for distal enhancer-promoter pairs, the condensate velocity *ν* is insufficient to bring the enhancer-promoter contact probability above the threshold. In these cases, cohesin is required to first bring the enhancer within a range where gradient-sensing by the condensate can bridge the remaining gap. These findings could explain recent experimental results that suggest transcription is dependent on cohesin for genomically distant enhancer-promoter pairs but independent for genomically proximal pairs^48,49^.

## V. CONDENSATES CAN EXHIBIT OSCILLATIONS IN MORPHOLOGY AND DISTANCE TO THE PROMOTER

Live-cell imaging experiments have observed condensates moving towards and away from promoters, resulting in dynamic “kissing” interactions^30,33^. In the cell, the transcription machinery enriched within transcriptional condensates first assembles into the preinitiation complex. RNA polymerase II then stalls near the promoter during promoter-proximal pausing, before finally proceeding to the productive elongation of RNA^58–60^. These processes give rise to an effective time delay between condensate-promoter contact and transcription. Hence, we next investigate whether such a delay could lead to oscillations in the condensate position relative to the promoter. We modify the reaction-diffusion model to incorporate a time delay *τ* into the transcription rate, and also to increase the sensitivity of the transcription rate to changes in the protein concentration,

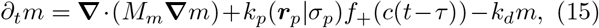

where *k*_*p*_(***r***_*p*_| *σ*_*p*_) is the Gaussian production rate that we previously used in Eq. (5). We introduce a Hill function *f*_+_(*c*) to increase the sensitivity of the transcription rate to changes in the protein concentration (Supplementary Information, Sec. SVII). In this model, proteins respond immediately to the RNA, whereas the RNA responds to the protein distribution from a previous time. We hypothesize that a delayed response of transcription to condensate-promoter contact, in combination with negative feedback between the protein and RNA, could lead to oscillations. In particular, accumulation of RNA at the promoter leads to charge repulsion, which could expel the condensate from the promoter.

To test these ideas, we reintroduce the re-entrant repulsion term, *γc*^2^*m*^2^, in the RNA-protein free energy [Eq. (2)] and simulate circular protein droplets using the same initial conditions as in Sec. III. When the condensate is far from the promoter, RNA concentrations are low and the attraction term *χcm* dominates [Eq. (2)], causing the condensate to move towards the promoter. When the condensate overlaps with the promoter, it enhances transcription after the time delay *τ*. As a result, the RNA concentration increases roughly 3.5-fold due to the increased sensitivity of the transcription rate to changes in protein concentration (Fig. 3A). At high RNA levels, the repulsion term *γc*^2^*m*^2^ dominates [Eq. (2)], expelling protein away from the promoter and returning the system to its basal rate of transcription (Fig. 3A).

**FIG. 3.**
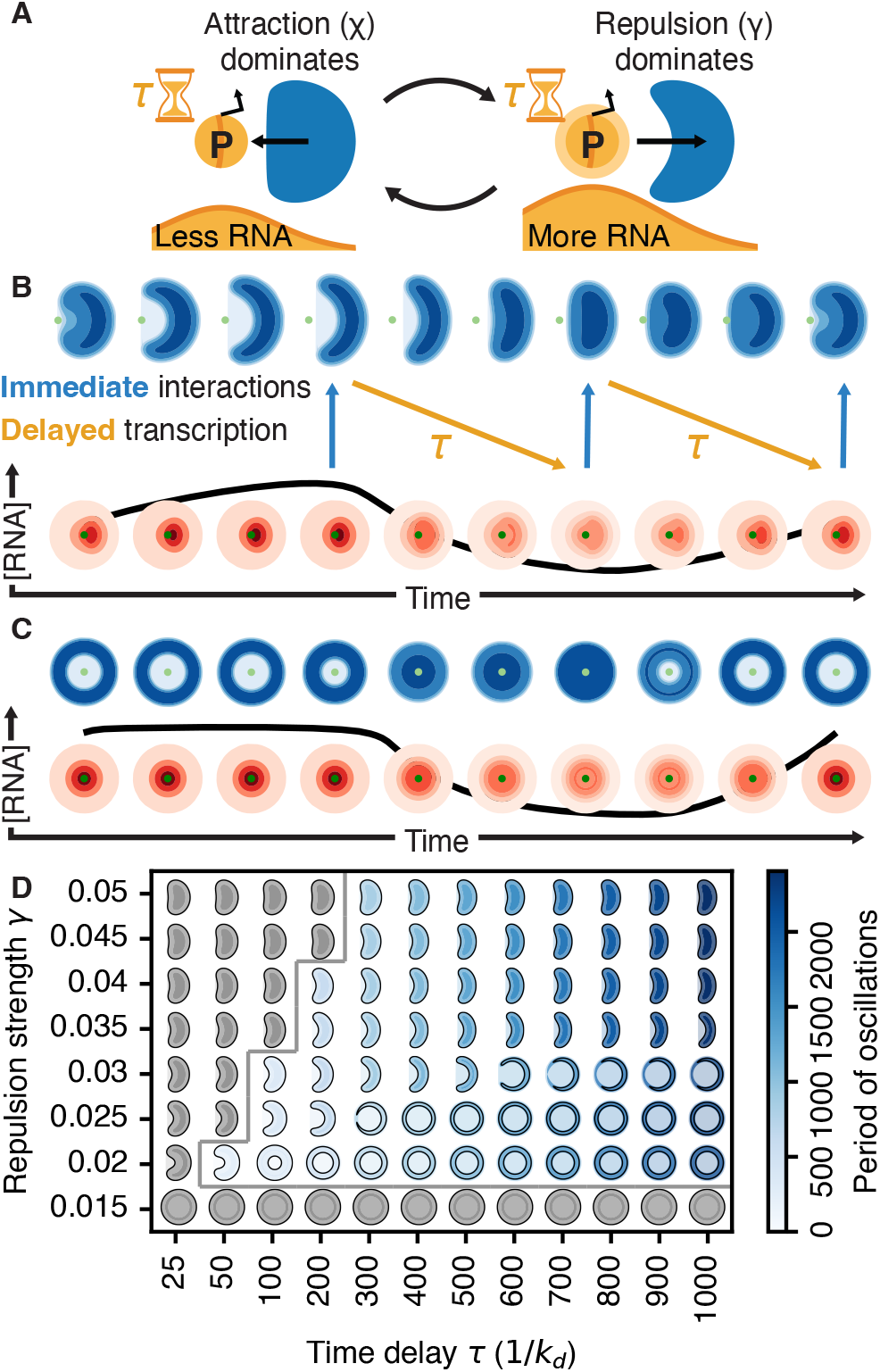
Condensate morphologies and RNA concentrations show oscillations when protein-RNA repulsion is sufficiently strong and transcription responds to the protein droplet after a delay time *τ*. (A) We consider a system where transcription at the promoter responds to the presence of protein after a fixed time delay. When the condensate overlaps with the promoter, increased transcription leads to an accumulation of RNA at the promoter. The resulting charge repulsion pushes the droplet away and deforms its shape. (B) Simulated trajectory of bean-like droplets (blue) and the RNA concentration field (red) over time with *γ* = 0.03 and *τ* = 250. The condensate morphology immediately responds to the concurrent RNA field, while transcription responds to a morphology from a previous time. (C) With *γ* = 0.02 and *τ* = 250, condensate morphologies oscillate between vacuoles and circular droplets (blue) in response to oscillating, radially symmetric RNA concentration fields (red). (D) Oscillations occur above a critical time delay and repulsion strength (indicated by the grey lines). The period of oscillations (colorbar) increases with the time delay. The repulsion strength determines the extent to which droplets colocalize with the promoter, which in turn determines the droplet morphology. Simulations parameters: *α* = 1, *β* = 0.25, *κ* = 0.1, *χ* = 0.1, *M*_*c*_ = 1, *D*_*m*_ = 1, *k*_*p*_ = 0.2, *σ*_*p*_ = 2.5, and *k*_*d*_ = 1.

The droplet exhibits two distinct types of oscillations. The droplet can either cycle through various bean-shaped morphologies (Fig. 3B) or alternate between a vacuole and a whole droplet (Fig. 3C). Which type of oscillation occurs depends on whether the repulsion strength and delay allow the droplet to fully colocalize with the promoter before RNA is enhanced by condensate-promoter overlap. When the droplet colocalizes with the promoter, the enhanced RNA levels push away the droplet symmetrically, forming a vacuole. When the droplet does not fully colocalize with the promoter, the enhanced RNA pushes it asymmetrically, forming a bean-shaped droplet. With strong repulsions *γ* ≥ 0.035, the droplet can never fully colocalize with the promoter even with an infinite time delay. Hence, the system oscillates between bean-like morphologies with different extents of condensate-promoter overlap (Fig. 3B). For weaker repulsions (*γ* = 0.02, 0.025, 0.03) the droplet can fully colocalize with the promoter given enough time, leading to symmetric oscillations centered at the origin, where a vacuole forms and refills (Fig. 3C). Reducing the time delay for these weaker repulsions leads to bean-like morphologies, as there is insufficient time for the droplet to flow and fully colocalize with the promoter at the origin.

The system exhibits oscillations only above a critical repulsion strength and time delay (Fig. 3D, grey line). The critical time delay is determined by the timescale for the condensate to flow towards the promoter from its initial position. This timescale increases as a function of the repulsion strength *γ*, since repulsion slows down the condensate velocity, as shown in our theoretical calculations (Supplementary Information, Sec. SIII C). There is also a critical repulsion strength below which oscillations do not occur. In this regime, the dynamics are dominated by the attractive protein-RNA interactions, even with transcription enhanced by the droplet, causing the droplet to reach a steady state where it simply colocalizes with the promoter.

The period of oscillations increases as ∽ 2*τ* (Fig. 3D, colorbar) and corresponds to ∼ 2*τ* plus an additional traversal time (Supplementary Information, Figs. S12 and S13). The droplet spends a time *τ* in both promoter-proximal and distal positions and requires some additional amount of time to traverse between these locations (Fig. 3B and C). The period of oscillations does not depend on the repulsion strength (Supplementary Information, Fig. S12). Overall, these results demonstrate that RNA concentrations and condensate morphologies can oscillate when transcription is delayed with respect to condensate-promoter contact. However, these results show oscillations in condensate morphology, which are different from the oscillations in condensate-promoter distance, or dynamic “kissing”, observed in live-cell imaging experiments^30,33^.

Therefore, we finally explore the conditions necessary for our system to exhibit center-of-mass oscillations. The droplet has no momentum under the overdamped dynamics of our system. Consequently, the center-of-mass displacement is solely determined by the range of RNA-driven repulsion, which can be extended by increasing the RNA diffusion length. We accomplish this by decreasing the degradation rate while maintaining the same concentration at the origin (Fig. 4A, B). When the diffusion length is larger than the size of the condensate, we find that the droplet can show center-of-mass oscillations (Fig. 4C). The center-of-mass oscillations are in phase with oscillations in the RNA concentration due to the delayed response of transcription to condensate-promoter contact (Fig. 4D). Our simulations are thus most consistent with a “hit and run” or “kiss and kick” picture of enhancer-promoter communication, where transcription is enhanced after the condensate has already moved away from the promoter^54,70^. Together, these results show that the combined effects of RNA-mediated directed motion, RNA-driven reentrant phase transitions, and time delays in gene bursting provide one possible explanation for the dynamic colocalization of transcriptional condensates with gene promoters.

**FIG. 4.**
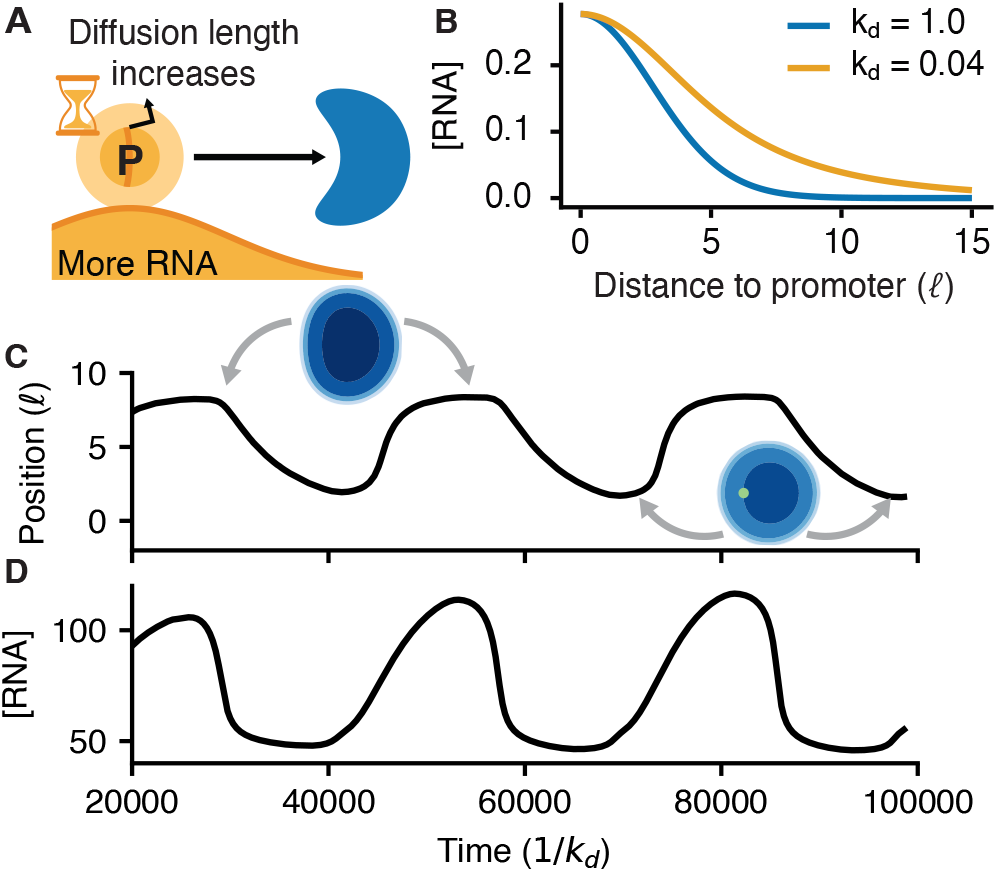
Condensates can show center-of-mass oscillations when the RNA diffusion length is larger than the condensate’s size. (A) Increasing the RNA diffusion length leads to a larger displacement of the droplet in response to RNA repulsion. (B) Decreasing the RNA degradation rate increases the diffusion length. We adjust the production rate such that the peak RNA concentration is unchanged. (C) Simulated trajectory shows the condensate moving towards and away from the promoter. The promoter (marked in green) is centered at the origin. (D) The RNA concentration is in phase with the condensate-promoter separation. Simulations parameters: *α* = 1, *β* = −0.25, *κ* = 0.1, *χ* = −0.1, *γ* = 0.06, *M*_*c*_ = 1, *D*_*m*_ = 1, *k*_*p*_ = 0.2, *σ*_*p*_ = 2.5, *k*_*d*_ = 0.04, and *τ* = 12000.

## VI. DISCUSSION

We have explored how protein condensates respond to an RNA transcription site (source) due to electrostatic interactions between proteins and RNA. Depending on the protein concentration and the RNA production rate, we found that the condensate can either move gradually toward the RNA source or dissolve and renucleate at the transcription site. Analytical calculations, in agreement with simulations, show that in the case of droplet motion, the droplet velocity is a non-monotonic function of the distance to the transcription site and peaks at a distance corresponding to the condensate radius. Interestingly, our calculations show that the liquid droplet, in *d* dimensions, moves *d* times faster than the individual proteins that make up the condensate. This enhancement in velocity is due to a *d* − 1-dimensional flux of material through the light phase that continuously nucleates the leading edge of the droplet as it moves^24^. In addition, material in the dense phase moves in the direction of the RNA gradient from the trailing edge to the leading edge^24,25^. Gene-specific parameters such as the transcription rate determine the steepness of the RNA concentration gradient, which in turn influences the speed of the droplet which senses this gradient.

In the present manuscript, condensate motion is driven by interactions with RNA in the bulk of the droplets and does not require hydrodynamic couplings *per se*. This mechanism stands in contrast with droplet propulsion due to the Marangoni effect^23^, and to the diffusiophoresis of colloids where solute interactions with the colloidal surface lead to tension gradients and a slip velocity against the solvent^71^. Although directed motion of transcriptional condensates has yet to be observed in experiments, the mechanism we identify is general for any droplet in a concentration gradient. For example, *in vitro*, it has been observed that salt concentration gradients induce phoretic condensate motion^39^. We speculate that this mechanism could contribute to the observed motion of nuclear speckles and Cajal bodies *in vivo*^34–36,72^. We next explored the consequences of directed motion of protein condensates towards actively transcribing promoters for enhancer-promoter communication. Since a transcriptional condensate is assumed to be bound to the enhancer locus^30^, its motion leads to a non-reciprocal attractive interaction between the enhancer and the promoter. Brownian dynamics simulations of the intervening chromatin chain show that directed motion increases the contact probability between the condensate and the promoter. This effect is more pronounced for genomically close enhancer-promoter pairs, since the condensate velocity is distance-dependent. In contrast, loop extrusion by cohesin should linearly compact the enhancer-promoter distance for any genomic separation between the enhancer and the promoter. We hypothesize that cohesin could reel in long-range enhancers while condensate directed motion could act at short and intermediate scales. Together, cohesin and condensates thus mediate a multi-scale enhancer-promoter attraction.

For genomically close enhancer-promoter pairs, condensate directed motion could provide a compensatory mechanism that allows for enhancer-promoter “microcompartments” (contacts) that are robust to cohesin depletion^46,51,52^. The increase in enhancer-promoter contact probability due to condensates alone could also explain why transcription appears to be cohesin-independent for genes with genomically close enhancers, but cohesin-dependent for genes with distal enhancers^48,49^. Consistent with our proposed mechanism, depletion of Mediator complex, which is a key component of transcriptional condensates, decreases enhancer-promoter interactions and associated gene expression levels^73^.

Future work could explore cohesin and condensate based mechanisms of gene regulation in more dynamic detail. For instance, the assumption that cohesin simply compacts the tether length connecting the enhancer and promoter fails to take into account the active dynamics of loop extrusion. Other works have suggested that loop extrusion could drag a promoter towards the surface of the condensate, leading to a nonmonotonic dependence of the enhancer-promoter contact probability on genomic distance^74^. In addition, in this work, we focus on steady state contact probabilities; however, the emergence of time-resolved live cell imaging data could enable direct visualization of the enhancer-promoter first-passage process^75–77^. Comparisons to such data would require simulating the active dynamics of loop extrusion, accounting for the finite lifetime of condensates and RNA gradients^30^, and calculating the first passage time to contact. Our polymer model could also be extended to couple the transcription rate at the promoter to the enhancer-promoter contact probability. Increased transcription would initially increase the velocity of enhancer-bound condensates and thus the enhancer-promoter contact probability, leading to a positive feed-back^42,78^. Eventually, accumulation of RNA could lead to charge repulsion which dissolves or expels the condensate, regulating the contact process in a complicated way. In this work, for simplicity, we have neglected the molecular interactions of individual proteins with chromatin^5,79^ and instead modeled the condensate as a single, large monomer of the chromatin polymer. Future work could incorporate the co-condensation of DNA with the proteins within the condensate, which increases the tension on the DNA chain^80^. Condensates have also been shown to mechanically exclude chromatin and pull together chromatin loci^81^. Other theoretical works have proposed that the polymer chain connecting the enhancer and promoter may be confined to the surface of the condensate^82^. There remains much to be learned about how condensates interact with DNA, indicating that further work is needed to determine the most appropriate approach for the multi-scale modeling of condensate-chromatin systems^83^.

Finally, we have shown that transcriptional condensates can show oscillations in morphology and in the distance to the promoter. Bean-shaped condensates and vacuoles have previously been reported as nonequilibrium steady states^27^ and observed in experiment^38^. We demonstrate that dynamic cycling between these states requires a time delay between condensate-promoter contact and transcription at the promoter. Such a delay is plausible biologically, due to the multiple internal state changes that the promoter undergoes prior to productive elongation of RNA. Together with negative feedback due to RNA accumulation at the promoter^12^, time delays can cause oscillations in condensate-promoter distance akin to “kissing” events^30,33^.

Unlike the observations in Ref.^30^, however, our model predicts that the RNA levels at the promoter are in phase with the condensate-promoter distance (out of phase with condensate-promoter contact). That is, transcription is enhanced after the condensate has moved away from the promoter, due to the time delay. Thus, our model provides a mechanistic basis for the proposed “hit and run” or “kiss and kick” model of enhancer-promoter communication^54,70^. More generally, our work demonstrates that phase separation of transcriptional proteins coupled with RNA production and degradation can yield rich dynamics with potentially far-reaching consequences for gene regulation.

## ACKNOWLEDGMENTS

The authors thank Mehran Kardar and Yannick Azhri Din Omar for their insightful discussions. We also thank Mehran Kardar for critical reading of the manuscript. We thank Thomas Nok Hin Cheng and the MIT Chemical Engineering Communication Lab for their feedback on figure presentation. This work was supported by the National Science Foundation, through the Biophysics of Nuclear Condensates grant (MCB-2044895). A.G. was also supported by an EMBO Postdoctoral Fellowship (ALTF 259-2022). D.K. was supported by the Graduate Research Fellowship Program under grant No. 2141064. A.K.C. and D.G. were also supported by the Ragon Institute. The authors acknowledge the MIT SuperCloud and Lincoln Laboratory Supercomputing Center for providing HPC resources that have contributed to the research results reported within this paper.

## References

1 S. F. Banani, H. O. Lee, A. A. Hyman, and M. K. Rosen, Nature reviews Molecular cell biology 18, 285 (2017).

2 Y. Shin and C. P. Brangwynne, Science 357, eaaf4382 (2017).

3 A. A. Hyman, C. A. Weber, and F. Jülicher, Annual Review of Cell and Developmental Biology 30, 39 (2014).

4 P. Li, S. Banjade, H. C. Cheng, S. Kim, B. Chen, L. Guo, M. Llaguno, J. V. Hollingsworth, D. S. King, S. F. Banani, P. S. Russo, Q. X. Jiang, B. T. Nixon, and M. K. Rosen, Nature 483, 336 (2012).

5 K. Shrinivas, B. R. Sabari, E. L. Coffey, I. A. Klein, A. Boija, A. V. Zamudio, J. Schuijers, N. M. Hannett, P. A. Sharp, R. A. Young, et al., Molecular cell 75, 549 (2019).

6 J. A. Morin, S. Wittmann, S. Choubey, A. Klosin, S. Golfier, A. A. Hyman, F. Jülicher, and S. W. Grill, Nature Physics 18, 271 (2022).

7 J. Berry, S. C. Weber, N. Vaidya, M. Haataja, C. P. Brangwynne, and D. A. Weitz, Proceedings of the National Academy of Sciences of the United States of America 112, E5237 (2015).

8 H. Zhang, S. Elbaum-Garfinkle, E. M. Langdon, N. Taylor, P. Occhipinti, A. A. Bridges, C. P. Brangwynne, and A. S. Gladfelter, Molecular Cell 60, 220 (2015).

9 M. Feric, N. Vaidya, T. S. Harmon, D. M. Mitrea, L. Zhu, T. M. Richardson, R. W. Kriwacki, R. V. Pappu, and C. P. Brangwynne, Cell 165, 1686 (2016).

10 P. R. Banerjee, A. N. Milin, M. M. Moosa, P. L. Onuchic, and A. A. Deniz, Angewandte Chemie - International Edition 56, 11354 (2017).

11 A. N. Milin and A. A. Deniz, Biochemistry 57, 2470 (2018).

12 J. E. Henninger, O. Oksuz, K. Shrinivas, I. Sagi, G. LeRoy, M. M. Zheng, J. O. Andrews, A. V. Zamudio, C. Lazaris, N. M. Hannett, et al., Cell 184, 207 (2021).

13 C. A. Weber, D. Zwicker, F. Jülicher, and C. F. Lee, Reports on Progress in Physics 82, 064601 (2019).

14 S. C. Glotzer, A. di Marzio Edmund, and M. Muthukumar, Phys. Rev. Lett. 73, 1919 (1994).

15 D. Carati and R. Lefever, Physical Review E 56, 3127 (1997).

16 J. D. Wurtz and C. F. Lee, Physical Review Letters 120, 078102 (2018).

17 J. Kirschbaum and D. Zwicker, Journal of The Royal Society Interface 18, 20210255 (2021).

18 Y. I. Li and M. E. Cates, Journal of Statistical Mechanics: Theory and Experiment 2020, 053206 (2020).

19 D. Zwicker, A. A. Hyman, and F. Jülicher, Physical Review E 92, 012317 (2015).

20 L. Demarchi, A. Goychuk, I. Maryshev, and E. Frey, Physical Review Letters 130, 128401 (2023).

21 S. F. Banani, A. Goychuk, P. Natarajan, M. M. Zheng, G. Dall’Agnese, J. E. Henninger, M. Kardar, R. A. Young, and A. K. Chakraborty, bioRxiv 10.1101/2024.10.12.614958 (2024).

22 D. Zwicker, R. Seyboldt, C. A. Weber, A. A. Hyman, and F. Jülicher, Nature Physics 13, 408 (2017).

23 S. Michelin, Annual Review of Fluid Mechanics 55, 77 (2023).

24 A. Goychuk, L. Demarchi, I. Maryshev, and E. Frey, Physical Review Research 6, 033082 (2024).

25 C. A. Weber, C. F. Lee, and F. Jülicher, New Journal of Physics 19, 053021 (2017).

26 P. C. Bressloff, Journal of Physics A: Mathematical and Theoretical 53, 365002 (2020).

27 H. H. Schede, P. Natarajan, A. K. Chakraborty, and K. Shrinivas, Nature Communications 14, 4152 (2023).

28 D. Hnisz, K. Shrinivas, R. A. Young, A. K. Chakraborty, and P. A. Sharp, Cell 169, 13 (2017).

29 B. R. Sabari, A. Dall’Agnese, A. Boija, I. A. Klein, E. L. Coffey, K. Shrinivas, B. J. Abraham, N. M. Hannett, A. V. Zamudio, J. C. Manteiga, et al., Science 361, eaar3958 (2018).

30 W.-K. Cho, J.-H. Spille, M. Hecht, C. Lee, C. Li, V. Grube, and I. I. Cisse, Science 361, 412 (2018).

31 S. Chong, C. Dugast-Darzacq, Z. Liu, P. Dong, G. M. Dailey, C. Cattoglio, A. Heckert, S. Banala, L. Lavis, X. Darzacq, and R. Tjian, Science 361, eaar2555 (2018).

32 A. Boija, I. A. Klein, B. R. Sabari, A. Dall’Agnese, E. L. Coffey, A. V. Zamudio, C. H. Li, K. Shrinivas, J. C. Manteiga, N. M. Hannett, et al., Cell 175, 1842 (2018).

33 M. Du, S. H. Stitzinger, J.-H. Spille, W.-K. Cho, C. Lee, M. Hijaz, A. Quintana, and I. I. Cissé, Cell 187, 331 (2024).

34 M. Platani, I. Goldberg, J. R. Swedlow, and A. I. Lamond, The Journal of cell biology 151, 1561 (2000).

35 M. Platani, I. Goldberg, A. I. Lamond, and J. R. Swedlow, Nature cell biology 4, 502 (2002).

36 S. M. Görisch, M. Wachsmuth, C. Ittrich, C. P. Bacher, K. Rippe, and P. Lichter, Proceedings of the National Academy of Sciences 101, 13221 (2004).

37 J. Kim, K. Y. Han, N. Khanna, T. Ha, and A. S. Belmont, Journal of cell science 132, jcs226563 (2019).

38 A. Pancholi, T. Klingberg, W. Zhang, R. Prizak, I. Mamontova, A. Noa, M. Sobucki, A. Y. Kobitski, G. U. Nienhaus, V. Zaburdaev, and L. Hilbert, Molecular Systems Biology 17, e10272 (2021).

39 V. S. Doan, I. Alshareedah, A. Singh, P. R. Banerjee, and S. Shin, Nature Communications 15, 7686 (2024).

40 E. Jambon-Puillet, A. Testa, C. Lorenz, R. W. Style, A. A. Rebane, and E. R. Dufresne, Nature Communications 15, 3919 (2024).

41 X. Wang, M. J. Cairns, and J. Yan, Nucleic Acids Research 47, 11481 (2019).

42 J. Zuin, G. Roth, Y. Zhan, J. Cramard, J. Redolfi, E. Piskadlo, P. Mach, M. Kryzhanovska, G. Tihanyi, H. Kohler, M. Eder, C. Leemans, B. van Steensel, P. Meister, S. Smallwood, and L. Giorgetti, Nature 604, 571 (2022).

43 J. H. Yang and A. S. Hansen, Nature Reviews Molecular Cell Biology 25, 574 (2024).

44 L. El Khattabi, H. Zhao, J. Kalchschmidt, N. Young, S. Jung, P. Van Blerkom, P. Kieffer-Kwon, K.-R. Kieffer-Kwon, S. Park, X. Wang, J. Krebs, S. Tripathi, N. Sakabe, D. R. Sobreira, S.-C. Huang, S. S. P. Rao, N. Pruett, D. Chauss, E. Sadler, A. Lopez, M. A. Nóbrega, E. L. Aiden, F. J. Asturias, and R. Casellas, Cell 178, 1145 (2019).

45 S. Cuartero, F. D. Weiss, G. Dharmalingam, Y. Guo, E. Ing-Simmons, S. Masella, I. Robles-Rebollo, X. Xiao, Y.-F. Wang, I. Barozzi, D. Djeghloul, M. T. Amano, H. Niskanen, E. Petretto, R. D. Dowell, K. Tachibana, M. U. Kaikkonen, K. A. Nasmyth, B. Lenhard, G. Natoli, A. G. Fisher, and M. Merkenschlager, Nature Immunology 19, 932 (2018).

46 M. J. Thiecke, G. Wutz, M. Muhar, W. Tang, S. Bevan, V. Malysheva, R. Stocsits, T. Neumann, J. Zuber, P. Fraser, S. Schoenfelder, J.-M. Peters, and M. Spivakov, Cell Reports 32, 107929 (2020).

47 L. Calderon, F. D. Weiss, J. A. Beagan, M. S. Oliveira, R. Georgieva, Y.-F. Wang, T. S. Carroll, G. Dharmalingam, W. Gong, K. Tossell, V. de Paola, C. Whilding, M. A. Ungless, A. G. Fisher, J. E. Phillips-Cremins, and M. Merkenschlager, eLife 11, e76539 (2022).

48 L. Kane, I. Williamson, I. M. Flyamer, Y. Kumar, R. E. Hill, L. A. Lettice, and W. A. Bickmore, Nature structural & molecular biology 29, 891 (2022).

49 N. J. Rinzema, K. Sofiadis, S. J. D. Tjalsma, M. J. A. M. Verstegen, Y. Oz, C. Valdes-Quezada, A.-K. Felder, T. Filipovska, S. van der Elst, Z. de Andrade dos Ramos, R. Han, P. H. L. Krijger, and W. de Laat, Nature Structural & Molecular Biology 29, 563 (2022).

50 T.-H. S. Hsieh, C. Cattoglio, E. Slobodyanyuk, A. S. Hansen, X. Darzacq, and R. Tjian, Nature Genetics 54, 1919 (2022).

51 A. Aljahani, P. Hua, M. A. Karpinska, K. Quililan, J. O. J. Davies, and A. M. Oudelaar, Nature Communications 13, 2139 (2022).

52 V. Y. Goel, M. K. Huseyin, and A. S. Hansen, Nature Genetics 55, 1048 (2023).

53 W. Bialek, T. Gregor, and G. Tka?cik, Action at a distance in transcriptional regulation (2019).

54 H. B. Brandão, M. Gabriele, and A. S. Hansen, Current Opinion in Cell Biology Cell Nucleus, 70, 18 (2021).

55 M.-T. Wei, Y.-C. Chang, S. F. Shimobayashi, Y. Shin, A. R. Strom, and C. P. Brangwynne, Nature Cell Biology 22, 1187 (2020).

56 K. Monfils and T. S. Barakat, European Journal of Cell Biology 100, 10.1016/j.ejcb.2021.151170 (2021).

57 M. Gabriele, H. B. Brandão, S. Grosse-Holz, A. Jha, G. M. Dailey, C. Cattoglio, T.-H. S. Hsieh, L. Mirny, C. Zechner, and A. S. Hansen, Science 376, 496 (2022).

58 J. Li, Y. Liu, H. S. Rhee, S. K. B. Ghosh, L. Bai, B. F. Pugh, and D. S. Gilmour, Molecular Cell 50, 711 (2013).

59 K. Adelman and J. T. Lis, Nature Reviews Genetics 13, 720 (2012).

60 L. Core and K. Adelman, Genes & Development 33, 960 (2019).

61 P. Natarajan, K. Shrinivas, and A. K. Chakraborty, Biophysical Journal 122, 2757 (2023).

62 M. Doi and S. F. Edwards, The Theory of Polymer Dynamics, International Series of Monographs on Physics Clarendon Press, Oxford, 2007).

63 V. I. Keizer, S. Grosse-Holz, M. Woringer, L. Zambon, K. Aizel, M. Bongaerts, F. Delille, L. Kolar-Znika, V. F. Scolari, S. Hoffmann, et al., Science 377, 489 (2022).

64 M. Socol, R. Wang, D. Jost, P. Carrivain, C. Vaillant, E. Le Cam, V. Dahirel, C. Normand, K. Bystricky, J.-M. Victor, et al., Nucleic acids research 47, 6195 (2019).

65 K. E. Polovnikov, H. B. Brandão, S. Belan, B. Slavov, M. Imakaev, and L. A. Mirny, Phys. Rev. X 13, 041029 (2023).

66 P. Mach, P. I. Kos, Y. Zhan, J. Cramard, S. Gaudin, J. Tünnermann, E. Marchi, J. Eglinger, J. Zuin, M. Kryzhanovska, et al., Nature Genetics 54, 1907 (2022).

67 B. Beltran, D. Kannan, Q. MacPherson, and A. J. Spakowitz, Physical Review Letters 123, 208103 (2019).

68 A. Goychuk, D. Kannan, A. K. Chakraborty, and M. Kardar, Proceedings of the National Academy of Sciences 120, e2221726120 (2023).

69 X. Darzacq, Y. Shav-Tal, V. De Turris, Y. Brody, S. M. Shenoy, R. D. Phair, and R. H. Singer, Nature structural & molecular biology 14, 796 (2007).

70 M. E. Pownall, L. Miao, C. E. Vejnar, O. M’Saad, A. Sherrard, M. A. Frederick, M. D. J. Benitez, C. W. Boswell, K. S. Zaret, J. Bewersdorf, and A. J. Giraldez, Science 381, 92 (2023).

71 S. Marbach, H. Yoshida, and L. Bocquet, Journal of fluid mechanics 892, A6 (2020).

72 T.-K. Kim, M. Hemberg, J. M. Gray, A. M. Costa, D. M. Bear, J. Wu, D. A. Harmin, M. Laptewicz, K. Barbara-Haley, S. Kuersten, E. Markenscoff-Papadimitriou, D. Kuhl, H. Bito, P. F. Worley, G. Kreiman, and M. E. Greenberg, Nature 465, 182 (2010).

73 S. Ramasamy, A. Aljahani, M. A. Karpinska, T. N. Cao, T. Velychko, J. N. Cruz, M. Lidschreiber, and A. M. Oudelaar, Nature Structural & Molecular Biology 30, 991 (2023).

74 T. Yamamoto, T. Sakaue, and H. Schiessel, Nucleic acids research 49, 5017 (2021).

75 H. Chen, M. Levo, L. Barinov, M. Fujioka, J. B. Jaynes, and T. Gregor, Nature genetics 50, 1296 (2018).

76 J. M. Alexander, J. Guan, B. Li, L. Maliskova, M. Song, Y. Shen, B. Huang, S. Lomvardas, and O. D. Weiner, eLife 10.7554/eLife.41769 (2019).

77 D. B. Brückner, H. Chen, L. Barinov, B. Zoller, and T. Gregor, Science 380, 1357 (2023).

78 J. Y. Xiao, A. Hafner, and A. N. Boettiger, eLife 10, e64320 (2021).

79 A. M. Chiariello, F. Corberi, and M. Salerno, Biophysical Journal 119, 873 (2020).

80 T. Quail, S. Golfier, M. Elsner, K. Ishihara, V. Murugesan, R. Renger, F. Jülicher, and J. Brugués, Nature Physics 17, 1007 (2021).

81 Y. Shin, Y.-C. Chang, D. S. Lee, J. Berry, D. W. Sanders, P. Ronceray, N. S. Wingreen, M. Haataja, and C. P. Brangwynne, Cell 175, 1481 (2018).

82 Q. Zhang, H. Shi, and Z. Zhang, A dynamic kissing model for enhancer-promoter communication on the surface of transcriptional condensate (2022).

83 M. Stortz, D. M. Presman, and V. Levi, Communications Biology 7, 1 (2024).

