## Supplementary Information for "RNA gradients can guide condensates toward promoters: implications for enhancer-promoter contacts and condensate-promoter kissing"

(Dated: November 4, 2024)

### CONTENTS

SI. Measurements from directed droplet motion 2

SII. Comparison of droplet velocity between simulation and theory 4

SIII. Derivation of droplet velocity in the sharp interface limit 6
A. Droplet velocity in 2D in response to a Gaussian RNA source 7
B. Droplet velocity in 3D in response to a point-like RNA source 8
C. Droplet velocity in response to arbitrary forcing 8

SIV. Measurements characterizing droplet oscillations 10

SV. Non-dimensionalization of model equations 16

SVI. Code availability 17

SVII. Increased sensitivity of transcription to proteins 18

SVIII. Units and conversions 19

SIX. Enhancer-Promoter contact probability is more sensitive to changes in condensate velocity for close pairs 20

SX. Effect of changing activity and friction of the enhancer 21

References 21

* These authors contributed equally to this work.
†
‡

### SI. MEASUREMENTS FROM DIRECTED DROPLET MOTION

In the present section, we discuss how we quantify the observables of the droplet, including its flow velocity towards the RNA source, using simulation data (Fig. S1). In our finite volume simulations, the concentration field is discretized into triangular mesh elements. We can extract from the simulations the concentration at each mesh element over time, the coordinates of the elements, and the area of these elements. We filter for the mesh elements with concentrations that exceed the critical point of the double well potential, $\bar{c}$, to define the droplet. This threshold, combined with the smooth interface of the droplet, can introduce "jitters" in the curves of the observables of the droplet. These jitters happen when a mesh element is included or excluded from the filter as the droplet interface evolves from one time step to the next. There is no noise in our deterministic simulations, and the jitters are just a limitation of how we discretize our simulations.

To calculate the flow velocity of the droplet, we need some way of quantifying the droplet's position. One way is to use the droplet's center of mass. We can calculate the center of mass by taking a weighted average of the position vectors within the droplet, $\mathcal{D}$, where the weights are given by the concentration field [Eq. S1]. Numerically, we calculate this weighted average using the concentration values ($c_i$), coordinates ($\boldsymbol{r}_i$), and area ($a_i$), at each of the droplet's mesh elements $i \in \mathcal{D}$:

$$\boldsymbol{r}_c = \frac{\int_{\mathcal{D}} \boldsymbol{r} c(\boldsymbol{r}) \, \mathrm{d}^2 r}{\int_{\mathcal{D}} c(\boldsymbol{r}) \, \mathrm{d}^2 r} \approx \frac{\sum_{i \in \mathcal{D}} \boldsymbol{r}_i c_i a_i}{\sum_{i \in \mathcal{D}} c_i a_i} \,. \tag{S1}$$

In our simulations, the condensate flows along the $x$-axis of the simulation domain because we initialize the droplet at position $(10, 0)$ and place the RNA source at position $(0, 0)$. It is the $x$-component of the center of mass vector $\boldsymbol{r}_c$ that we show in Fig. S1A.

We calculate the droplet velocity by taking the numerical derivative of the center of mass over the simulation time using NumPy's `numpy.gradient` function [1]. Note that this gradient calculation accentuates tiny jitters in the center of mass, which are introduced by our filtering (Fig. S1B, Raw).

To plot a smoother trajectory, we use the moving average of the center of mass over time, which we get using SciPy's `scipy.ndimage.uniform_filter1d` function (Fig. S1B, Moving average) [2]. In the beginning of the simulation, there are large fluctuations in the velocity, which is likely a numerical artifact (Fig. S1B) This artifact may be caused by the droplet's initial expansion from its initial conditions. The droplet initially expands because we nucleate a sharp circular patch with a concentration that is higher than the dense phase concentration. This effect may also be accentuated by dividing over the smaller time steps from the first few iterations of our adaptive time-stepping scheme when calculating the velocity.

To better understand how the simulation of the full dynamics and the analytical theory for sharp and round droplets deviate from one another, we looked into how various terms change over time. The droplet area (Fig. S1C) and the difference between the dense and light phase concentrations (Fig. S1D,E) both enter into the expression for the flow velocity. The droplet grows in size as it approaches the RNA source because the attractive potential ($\chi m$) increases with RNA concentration, recruiting more proteins into the droplet. The dense and light phase concentrations are not exactly at the binodal points. The elongation of the droplet, which can be measured by the eccentricity (Fig. S1F), quantifies how the round droplet assumption in the theory breaks down in the parameter regime of our simulations. With the full dynamics, the droplet can undergo significant elongation because the RNA gradients at the front and back interfaces of the droplet are different. Another factor which can cause differences in dynamics between simulation and theory, which we do not quantify, is the finite interface width of the simulated droplets, which differs from the sharp interface assumed in the theory.

We calculate the area of the droplet by summing the areas of the mesh elements within the droplet. We determine the concentrations of the dense and light phases by calculating the weighted average of the concentrations within ($\mathcal{D}$) and outside ($\mathcal{D}^c$) the droplet, respectively, using the areas of the mesh elements as weights:

$$c_+ = \frac{\int_{\mathcal{D}} c(\boldsymbol{r}) \mathrm{d}^2 r}{\int_{\mathcal{D}} \mathrm{d}^2 r} \approx \frac{\sum_{i \in \mathcal{D}} c_i a_i}{\sum_{i \in \mathcal{D}} a_i} \,, \tag{S2}$$

$$c_- = \frac{\int_{\mathcal{D}^c} c(\boldsymbol{r}) \mathrm{d}^2 r}{\int_{\mathcal{D}^c} \mathrm{d}^2 r} \approx \frac{\sum_{i \notin \mathcal{D}} c_i a_i}{\sum_{i \notin \mathcal{D}} a_i} \,. \tag{S3}$$

We calculate the aspect ratio using the width and height of the droplet, which we obtain using the differences in the maximum and minimum coordinates of the droplet's mesh elements in the axis of flow and the perpendicular axis. For similar reasons to that of the velocity, this calculation introduces jitters when mesh elements are included and excluded from the droplet as the interface evolves.

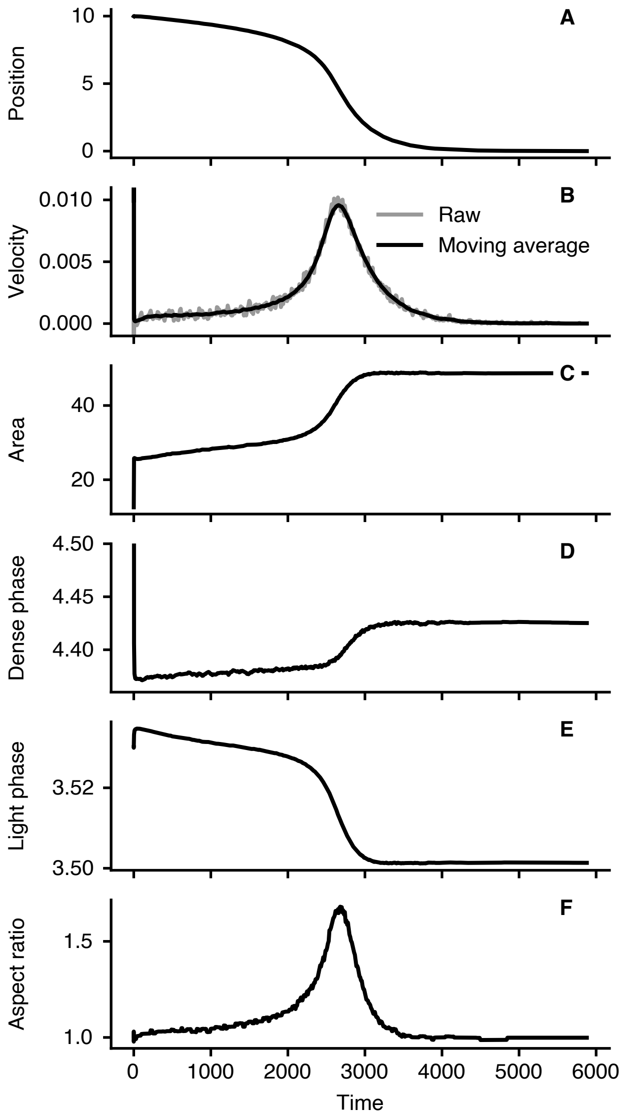


FIG. S1. Quantification of droplet properties from 2D simulations during flow towards a Gaussian source. (A) Center of mass of the droplet in the axis of flow. (B) Velocity of the droplet in the direction of flow. (C) Area of the droplet. (D) Dense phase (condensate) concentration. Initial condition is $c_{+}(t = 0) = 5.5$. (E) Light phase (background) concentration. (F) Aspect ratio (elongation) of the droplet.

### SII. COMPARISON OF DROPLET VELOCITY BETWEEN SIMULATION AND THEORY

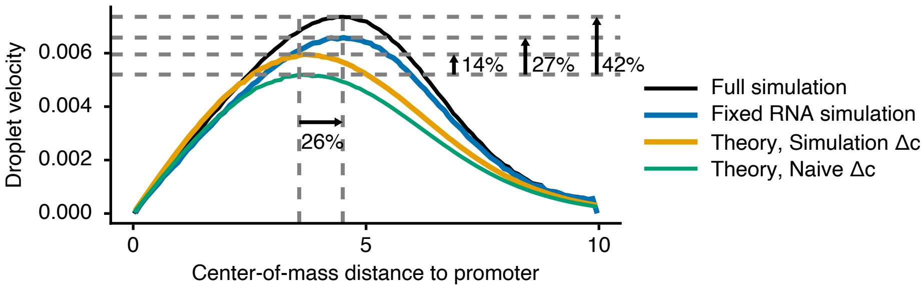


FIG. S2. Droplet velocity as a function of the distance between the droplet and the promoter. Full simulation (black line): velocity calculated from a simulation of Eqs. (4) and (5) in the main text. Fixed RNA simulation (blue line): velocity calculated from a simulation with the RNA production rate fixed to be $k_p c_-$ so that the RNA field does not respond to the droplet. Theory, Simulation $\Delta c$ (yellow line): velocity calculated from Eq. (S19) using the droplet volume ($V_\mathcal{D}$), dense phase concentration ($c_+$), and light phase concentration ($c_-$) from simulation data. Theory, Naive $\Delta c$ (green line): velocity calculated from Eq. (S19) using the droplet volume ($V_\mathcal{D}$) from simulation data, but assuming dense phase concentration $c_+ = 4.5$ and light phase concentration $c_- = 3.5$. The differences between the magnitude of the simulation and theory curves decrease when we account for the response of the RNA field to the moving droplet and the change in $\Delta c$ during droplet motion.

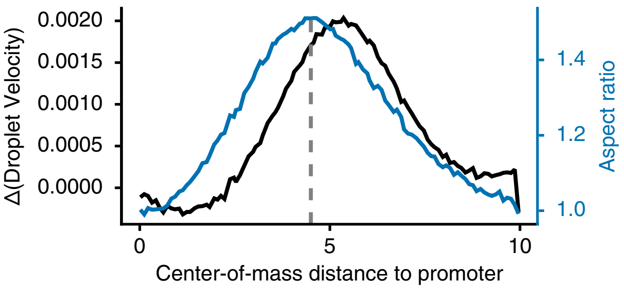


FIG. S3. Difference in droplet velocity from the simulation and theory as a function of the distance between the droplet and the promoter. Refer to Fig. S2 for a description of the simulations and calculations using the theory.

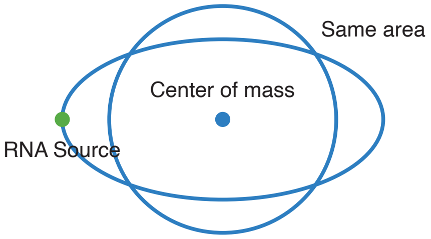


FIG. S4. An elongated droplet with the same area and center-of-mass distance as a circular droplet. The elongated droplet reaches the RNA source whereas the circular droplet does not. Hence, the peak velocity in simulations occurs at a larger distance than in theory.

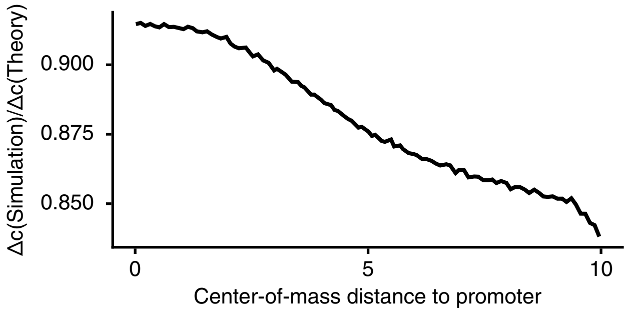


FIG. S5. Fold change of the difference in average protein concentrations within and outside the droplet from simulations relative to the difference between binodal points ($c_+ = 4.5$ and $c_- = 3.5$), as a function of the distance between the droplet and the promoter.

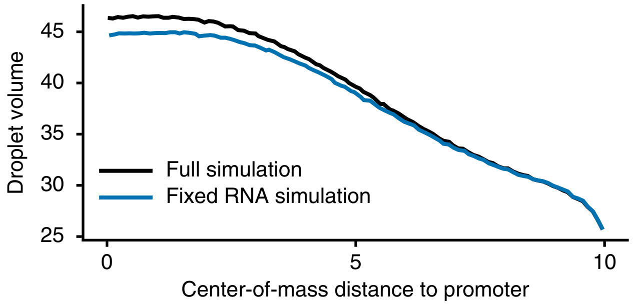


FIG. S6. Volume of the droplet as a function of the distance between the droplet and the promoter. Refer to Fig. S2 for a description of the simulations and calculations using the theory.

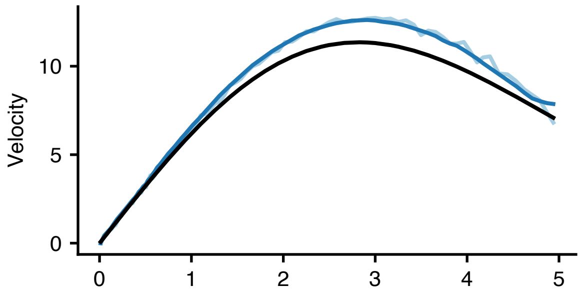


FIG. S7. Droplet velocity as a function of distance to the promoter for a round droplet.

### SIII. DERIVATION OF DROPLET VELOCITY IN THE SHARP INTERFACE LIMIT

To analyze droplet flow analytically, we seek a traveling-wave solution $c(\boldsymbol{x}-\boldsymbol{v}t) \equiv c(\boldsymbol{z})$ to the continuity equation for the protein dynamics. Such an ansatz results in a steady state in the co-moving frame,

$$-\boldsymbol{v}\cdot\boldsymbol{\nabla}c(\boldsymbol{z}) = \boldsymbol{\nabla}\cdot[M_c(\boldsymbol{\nabla}\mu_{cc}+\boldsymbol{\nabla}\mu_{cm})]. \tag{S4}$$

We next invoke the sharp interface approximation, which assumes that the interface width $w = \sqrt{2\kappa/\beta}$ is smaller than all other length scales relevant to the dynamics. Mathematically, we simultaneously take the limits $\alpha \to \infty$ and $\beta \to \infty$, such that the light and dense phase protein concentrations ($c^*_\pm = \bar{c} \pm \sqrt{\beta/\alpha}$) remain finite. Note that in the Cahn-Hilliard model, the effective surface tension of the droplet [3],

$$\gamma = \frac{1}{6}(\Delta c)^2\sqrt{2\kappa\beta}, \tag{S5}$$

diverges when one takes the above limit, implying a round droplet shape [4]. We thus consider a piecewise constant protein concentration profile $c(\boldsymbol{z})$ which adopts the dense phase equilibrium value $c^*_+$ inside the droplet and the light phase equilibrium concentration $c^*_-$ outside the droplet. In order to analyze (S4) in the sharp interface limit, where gradients become singular at the droplet interface, following Ref. [4], we re-arrange (S4) as

$$\nabla\cdot\boldsymbol{J}_\Delta = 0, \tag{S6}$$

where the current can be viewed as an integral of (S4),

$$\boldsymbol{J}_\Delta = M_c(\boldsymbol{\nabla}\mu_{cc}+\boldsymbol{\nabla}\mu_{cm}) + \boldsymbol{v}(c(\boldsymbol{z})-c_-). \tag{S7}$$

The integration constant $\boldsymbol{v}c_-$ ensures that the current vanishes in the far-field ($|\boldsymbol{z}| \to \infty$), where all concentration gradients vanish and the protein concentration is given by the light phase concentration $c_-$.

After imposing a piecewise constant protein concentration profile, Eq. (S4) has two unknowns, the droplet velocity $\boldsymbol{v}$ and the function $\mu_{cc}$. In order to solve for the velocity, we therefore need to impose a constraint on $\mu_{cc}(c(\boldsymbol{z})|\boldsymbol{v})$ to ensure thermodynamic consistency of the traveling wave ansatz. In Eq. (S4), the effective force due to protein-RNA interactions, $-\nabla\mu_{cm}$, breaks energy conservation because RNA is continuously produced and degraded. In contrast, the current of proteins $\boldsymbol{j} = c(\boldsymbol{z})\boldsymbol{v}$ in the presence of the purely thermodynamic force $-\nabla\mu_{cc}$ should not dissipate power [4],

$$\partial_t\mathcal{F}_{cc}[c] = \boldsymbol{v}\cdot\int \mathrm{d}^d\boldsymbol{z}\;\; c(\boldsymbol{z})\boldsymbol{\nabla}\mu_{cc} = 0. \tag{S8}$$

Because $\mu_{cc}$ far from the droplet is uniform, following Ref. [4], one can show that Eq. (S8) reduces to the following thermodynamic consistency condition involving an integral over the domain $\mathcal{D}$ of the droplet, $|\boldsymbol{z}| \leq R$,

$$\int_{\mathcal{D}} \mathrm{d}^d\boldsymbol{r}\;\; \hat{\boldsymbol{e}}_v\cdot\boldsymbol{\nabla}\mu_{cc} = 0. \tag{S9}$$

The condition in Eq. (S9) implies that the effective force due to protein-protein interactions alone must be symmetric about the droplet center in the direction of flow $\hat{\boldsymbol{e}}_v$. If Eq. (S9) does not hold, then the net force due to protein-protein interactions alone would cause the droplet to accelerate or decelerate in the absence of an RNA field. Together, Eqs. (S6) and (S9) can be solved for $\mu_{cc}$ and the droplet velocity.

First, we substitute $\mu_{cm} = \delta\mathcal{F}_{cm}/\delta c = \chi\nabla m$ into Eq. (S4) [cf. also the integration constant in Eq. (S7)], where $m(\boldsymbol{z})$ is the steady state RNA concentration profile in the co-moving frame and $\chi$ is the protein-RNA interaction parameter,

$$M_c(\nabla^2\mu_{cc}(\boldsymbol{z}) + \chi\nabla^2 m(\boldsymbol{z})) + \boldsymbol{v}\cdot\nabla(c(\boldsymbol{z})-c_-) = 0. \tag{S10}$$

Performing a Fourier transform on all the fields, $\mu \to \tilde{\mu}_q$, $m \to \tilde{m}_q$, and $c \to \tilde{c}_q$ and solving for $\mu_{cc}$ gives

$$\mu_{cc}(\boldsymbol{z}) = \int \frac{d^d\boldsymbol{q}}{(2\pi)^d}\left[\frac{i\boldsymbol{v}\cdot\boldsymbol{q}}{M_c q^2}c_q - \chi m_q\right]e^{i\boldsymbol{q}\cdot\boldsymbol{z}}, \tag{S11}$$

where

$$c_q = \int d^d\boldsymbol{z}(c(\boldsymbol{z}) - c_-)e^{-i\boldsymbol{q}\cdot\boldsymbol{z}} = \Delta c \int_{\mathcal{D}} d^d\boldsymbol{z} e^{-i\boldsymbol{q}\cdot\boldsymbol{z}} := \Delta c g(-\boldsymbol{q}). \tag{S12}$$

Here, we have defined $\Delta c = c_+^* - c_-^*$ as the difference between the dense and light phase concentrations, and $\mathcal{D}$ indicates the domain of the droplet volume. If we then apply the condition in Eq. (S9), we find

$$\int_{\mathcal{D}} \mathrm{d}^d\boldsymbol{z}\ \ \hat{\boldsymbol{e}}_v \cdot \boldsymbol{\nabla}\mu_{cc} = -\int_{\mathcal{D}} d^d\boldsymbol{z} \left\{ \hat{\boldsymbol{e}}_v \cdot \left[ \frac{v\Delta c}{M_c} \int \frac{d^d\boldsymbol{q}}{(2\pi)^d} \left( \boldsymbol{q} \frac{\hat{\boldsymbol{e}}_v \cdot \boldsymbol{q}}{q^2} g(-\boldsymbol{q}) e^{i\boldsymbol{q}\cdot\boldsymbol{z}} \right) + \chi \boldsymbol{\nabla} m(\boldsymbol{z}) \right] \right\} = 0. \tag{S13}$$

Exchanging the order of integration in the first term and substituting $g(\boldsymbol{q}) = \int_{\mathcal{D}} d^d\boldsymbol{z} e^{i\boldsymbol{q}\cdot\boldsymbol{z}}$ gives the following solution for the droplet velocity in $d$ dimensions:

$$\frac{v\Delta c}{M_c} \int \frac{d^d\boldsymbol{q}}{(2\pi)^d} (\hat{\boldsymbol{e}}_v \cdot \hat{\boldsymbol{q}})^2 |g(\boldsymbol{q})|^2 = -\chi \int_{\mathcal{D}} \mathrm{d}^d\boldsymbol{z}\ \hat{\boldsymbol{e}}_v \cdot \boldsymbol{\nabla} m(\boldsymbol{z}). \tag{S14}$$

The right-hand side demonstrates that droplet flow is driven by the gradient in the RNA concentration field and is proportional to the strength of protein-RNA interactions, $\chi$. When $\chi < 0$ (attractive protein-RNA interactions), as considered in the main text, droplet flow is in the direction of the RNA gradient. When $\chi > 0$ (repulsive interactions), the droplet flows against the gradient.

To obtain the velocity as a function of distance to the promoter, $r$, we need to compute the steady-state RNA concentration field $m(\boldsymbol{z}|r)$. Relative to the droplet center of mass, RNA is produced at a location $\boldsymbol{r}_p = r\hat{\boldsymbol{e}}_v$. We further assume that the production rate $k_p(\boldsymbol{z}|\boldsymbol{r}_p)c_-$ is purely determined by the protein concentration in the light phase (i.e. unaffected by the moving droplet). Under these assumptions, RNA production is balanced by degradation and diffusion, resulting in the steady state,

$$D_m \nabla^2 m + k_p(\boldsymbol{z}|r\hat{\boldsymbol{e}}_v)c_- - k_d m = 0. \tag{S15}$$

Thus, for each value of the condensate-promoter distance $r$, we compute the solution to the inhomogeneous Helmholtz equation, which is a convolution of the production rate and the Green's function,

$$m(\boldsymbol{z}|r) = \frac{c_-}{D_m} \int d^d\boldsymbol{z}' G(\boldsymbol{z} - \boldsymbol{z}') k_p(\boldsymbol{z}'|r\hat{\boldsymbol{e}}_v), \tag{S16}$$

and substitute into Eq. (S14).

In order to perform the integral on the left-hand side of Eq. (S14), we must specify a coordinate system and a direction of flow. Below, we carry out the calculations in $d = 2$ and $d = 3$ dimensions using polar and spherical coordinates, respectively.

#### A. Droplet velocity in 2D in response to a Gaussian RNA source

Without loss of generality, assume the circular droplet moves along the $\hat{\boldsymbol{e}}_v = \hat{\boldsymbol{x}}$ direction. Then, in polar coordinates,

$$g(\boldsymbol{q}) = \int_0^{2\pi} d\phi \int_0^R dr\ \ r e^{iqr\cos\phi} = \frac{2\pi R}{|\boldsymbol{q}|} J_1(R|\boldsymbol{q}|), \tag{S17}$$

where $R$ is the radius of the droplet and $J_1(x)$ is a Bessel function of the first kind. The left-hand side of Eq. (S14) is then

$$\frac{v\Delta c}{4\pi^2 M_c} \int_0^{2\pi} d\phi \cos^2\phi \int_0^{\infty} dq\ \ q\,|g(\boldsymbol{q})|^2 = \frac{v\Delta c}{2M_c}(\pi R^2). \tag{S18}$$

Thus, the velocity of a 2D circular droplet in response to an RNA gradient is given by

$$v = -\frac{2M_c\chi}{\Delta c\,(\pi R^2)} \int_{r\le R} d^2\boldsymbol{r}\ \ \hat{x} \cdot \boldsymbol{\nabla} m(\boldsymbol{r}) = -\frac{2M_c\chi}{\Delta c(\pi R^2)} \int_0^{2\pi} R d\phi\ \ m(R,\phi)\cos\phi, \tag{S19}$$

where in the final equality we have applied the divergence theorem.

The solution to the 2D Helmholtz equation, Eq. (S16), with a Gaussian production rate is given by

$$m(\boldsymbol{z}) = \frac{k_p c_-}{2\pi D_m} \int d^2\boldsymbol{z}' K_0\left(\frac{|\boldsymbol{z}-\boldsymbol{z}'|}{\ell}\right) \exp\left(-\frac{(\boldsymbol{z}'-r\hat{\boldsymbol{x}})^2}{2\sigma_p^2}\right), \tag{S20}$$

where the Green's kernel is a modified Bessel function of the second kind, and $\ell = \sqrt{D_m/k_d}$ is the RNA diffusion length. Since Eq. (S20) does not have a closed form expression, we numerically integrate Eq. (S19) to obtain the droplet velocity as a function of distance to the Gaussian source. In the absence of an analytical expression for the droplet radius $R$ as a function of time, we take the droplet radius from 2D simulation data.

#### B. Droplet velocity in 3D in response to a point-like RNA source

Next, we consider spherical droplets in 3D which flow in the $\hat{\boldsymbol{z}}$ direction. In spherical coordinates, the left-hand side of Eq. (S14) is

$$\frac{v\Delta c}{(2\pi)^2 M_c} \int_0^\pi d\theta \sin\theta \cos^2\theta \int_0^\infty dq \;\; q^2 |g(\boldsymbol{q})|^2 = \frac{v\Delta c}{3M_c}\left(\frac{4}{3}\pi R^3\right). \tag{S21}$$

Thus, the velocity of a 3D spherical droplet in response to an RNA gradient is given by

$$v = -\frac{3M_c\chi}{\Delta c(4\pi R^3/3)} \int_{r\leq R} d^3\boldsymbol{r} \;\; \partial_z m(\boldsymbol{r}) = -\frac{3M_c\chi}{\Delta c(4\pi R^3/3)} \int_0^{2\pi} d\phi \int_0^\pi d\theta R^2 \sin\theta \;\; m(R,\theta,\phi)\cos\theta, \tag{S22}$$

where in the final equality we have applied the divergence theorem. Comparing Eq. (S19) and Eq. (S22), the numerical prefactor for the droplet velocity is simply the dimension $d$. For an RNA point source located at $z_0\hat{\boldsymbol{z}}$ relative to the droplet, the solution to the 3D Helmholtz equation, (S16), with $k_p(\boldsymbol{r}|z_0) = k_p\delta^3(\boldsymbol{r} - z_0\hat{\boldsymbol{z}})$, is

$$m(\boldsymbol{r}) = \frac{k_p c_-}{4\pi D} \frac{\mathrm{e}^{-|\boldsymbol{r}-z_0\hat{\boldsymbol{z}}|/\ell}}{|\boldsymbol{r}-z_0\hat{\boldsymbol{z}}|}. \tag{S23}$$

Substituting Eq. (S23) into Eq. (S22) gives

$$v = -\frac{3M_c\chi k_p c_-}{2\Delta c(4\pi R^3/3)D_m} \frac{l}{z_0^2}\left[\mathrm{e}^{-\frac{R+z_0}{\ell}}(\ell+R)(\ell+z_0) \; - \mathrm{e}^{-\frac{|R-z_0|}{\ell}}(\ell^2 - Rz_0 + \ell|R-z_0|)\right]. \tag{S24}$$

To gain insight into this expression, we consider the limit of infinite diffusion length $\ell \to \infty$ and find that the velocity is linear in $z_0$ and inversely proportional to the droplet volume for $z_0 < R$. The velocity is sharply peaked at $z_0 = R$ and then falls off as $z_0^{-2}$ for $z_0 > R$ independent of the droplet radius. The large $z_0$ behavior shows that away from the point source the droplet is simply sensing gradients in $m$ which fall off as $z_0^{-2}$ [cf. Eq. (S23)].

#### C. Droplet velocity in response to arbitrary forcing

The above derivation can be generalized to describe steady motion of a droplet in response to arbitrary forcing, $\boldsymbol{f}$,

$$-\boldsymbol{v}\cdot\boldsymbol{\nabla}c(\boldsymbol{z}) = \boldsymbol{\nabla}\cdot[M_c(\boldsymbol{\nabla}\mu_{cc} - \boldsymbol{f})]. \tag{S25}$$

In $d = 2$ or $d = 3$ dimensions, the velocity of the droplet is then given by

$$v = \frac{dM_c}{\Delta c V_{\mathcal{D}}} \int_{\mathcal{D}} d^3\boldsymbol{z} \;\; (\hat{\boldsymbol{e}}_v \cdot \boldsymbol{f}), \tag{S26}$$

where in the present work we consider $\boldsymbol{f} = -\boldsymbol{\nabla}\mu_{cm} = -\chi\boldsymbol{\nabla}m - \gamma m^2\boldsymbol{\nabla}c - 2\gamma cm\boldsymbol{\nabla}m$ [cf. Eq. (2) in the main text]. Thus, for attractive protein-RNA interactions at low RNA concentration ($\chi < 0$) and repulsive protein-RNA interactions at high RNA concentration ($\gamma > 0$), the velocity of the droplet,

$$v = \frac{dM_c}{\Delta c V_{\mathcal{D}}} \int_{\mathcal{D}} d^3\boldsymbol{z} \;\; [\hat{\boldsymbol{e}}_v \cdot (|\chi| - 2|\gamma|c_+ m)\boldsymbol{\nabla}m], \tag{S27}$$

slows in the presence of repulsive interactions, and can even reverse direction.

### SIV. MEASUREMENTS CHARACTERIZING DROPLET OSCILLATIONS

In the present section, we discuss how we quantify the period of the oscillations. We found that the droplet's properties outlined in Fig. S1 cannot always be used to quantify oscillations: with certain parameters, the droplet undergoes donut-like (vacuole-forming) oscillations where the center of mass and other properties do not change with time (Figs. S9A, C, and D). We therefore calculate the total amount of RNA in the system (Eq. S28), which must oscillate for the droplet to oscillate as well:

$$m_{\text{total}} = \int m(\boldsymbol{r})\mathrm{d}^2 r \approx \sum_{\forall i} m_i a_i \,. \tag{S28}$$

The RNA production term, and hence the amount of RNA, must both oscillate because this is where we introduced the time delay [Eq. (15)]. We use `scipy.signal.find_peaks` on the RNA amount over time to get the peaks and troughs in RNA (Figs. S8B and S9B). Using these points, we calculate, at steady state, the period of the oscillations and the change in RNA amount. We also use the same method to calculate the center of mass displacement during oscillations for bean-like (asymmetric) oscillations.

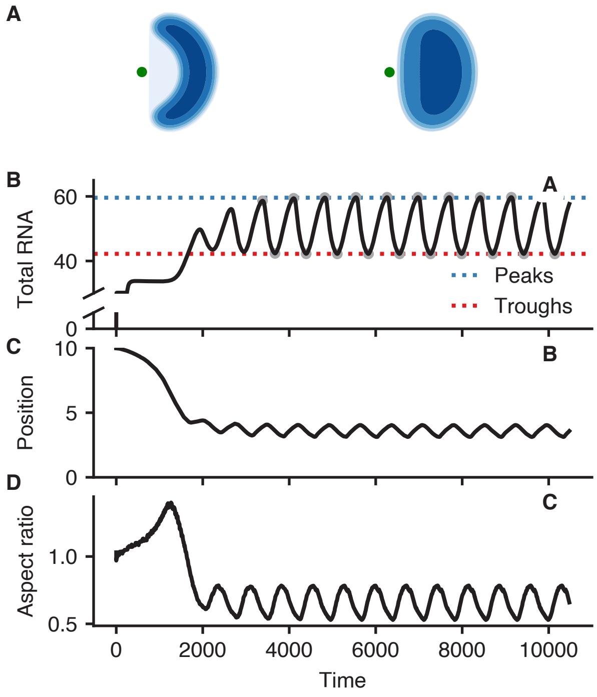


FIG. S8. Quantification of bean-like (asymmetric) oscillations. (A) Bean-like morphologies at peak and trough RNA production. This phenomenology occurs when the droplet is prevented from travelling to the center of the promoter, which happens when the repulsion strength is strong enough or the time delay is not long enough. The droplet retains memory of its initial condition. (B) Oscillations in RNA amount occur as the protein is attracted towards and repelled away from the promoter. We quantify the period from the oscillations in the RNA amount as it is a direct effect of the RNA production term's delayed response to the protein. (C) Short-distance center of mass oscillations. (D) Aspect ratio changes with time because the bean-like morphologies are dynamic.

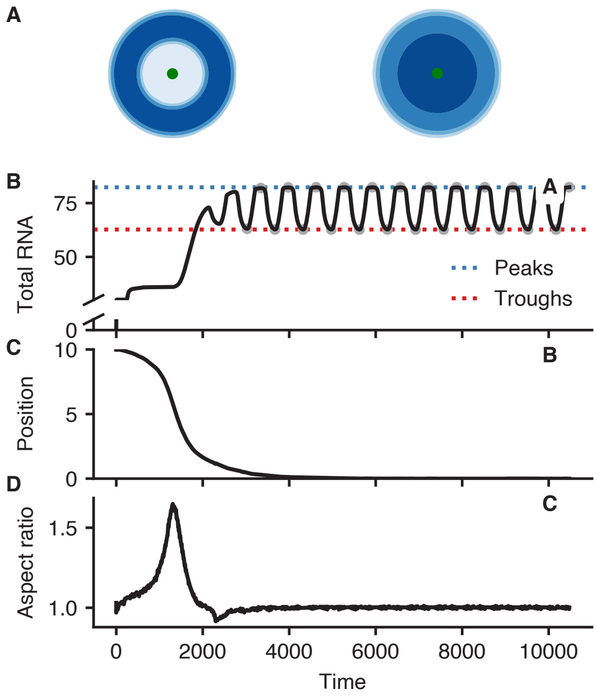


FIG. S9. Quantification of donut-like (vacuole-forming) oscillations. (A) Donut-like morphologies at peak and trough RNA production. This phenomenology occurs when the droplet travels to the center of the promoter, which happens when the repulsion strength is weak enough or the time delay is long enough. The droplet does not retain memory of its initial condition. (B) Oscillations in RNA amount occur as the protein is attracted towards and repelled away from the promoter. (C) No center of mass oscillations occur because the droplet remains centered at the promoter and the repulsions do not break apart the droplet. (D) Aspect ratio remains at one because the droplet remains spherical.

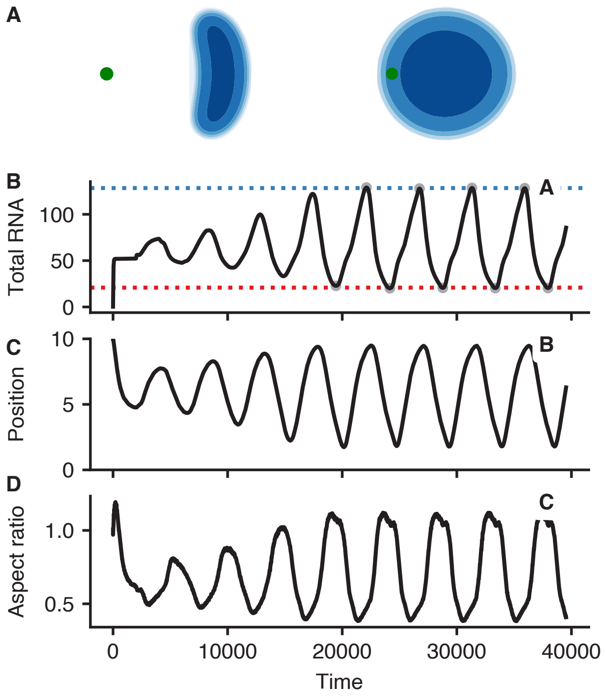


FIG. S10. Quantification of bean-like oscillations with large displacements. Simulations are under a different parameter regime of higher diffusion length, and attraction and repulsion increased by the same factor. (A) In a different parameter regime, the condensate can be repelled further away from the promoter. (B) Oscillations in RNA amount are larger because the protein is repelled further away from the promoter. (C) Condensate experiences center-of-mass displacements in the same order of magnitude as its size. (D) Oscillations in aspect ratio are larger because the condensate elongates as it flows back to the promoter.

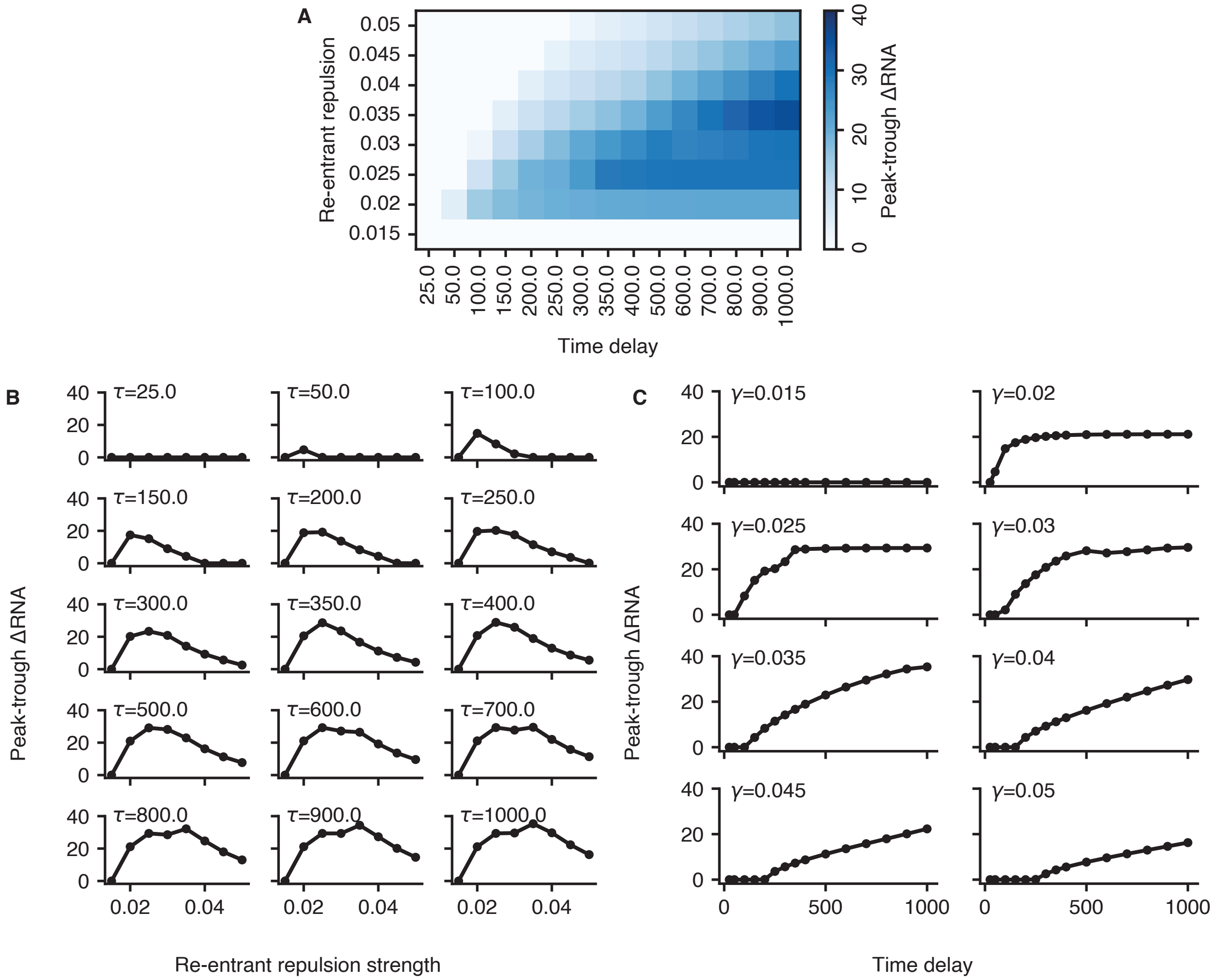


FIG. S11. (A) Change in peak and trough RNA amount depends on the time delay ($\tau$) and the strength of protein-RNA repulsion at high RNA concentrations ($\gamma$). (B) Change in peak and trough RNA amount as a function of repulsion strength, for fixed time delays $\tau$. (C) Change in peak and trough RNA amount as a function of time delay, for fixed repulsion strength $\gamma$.

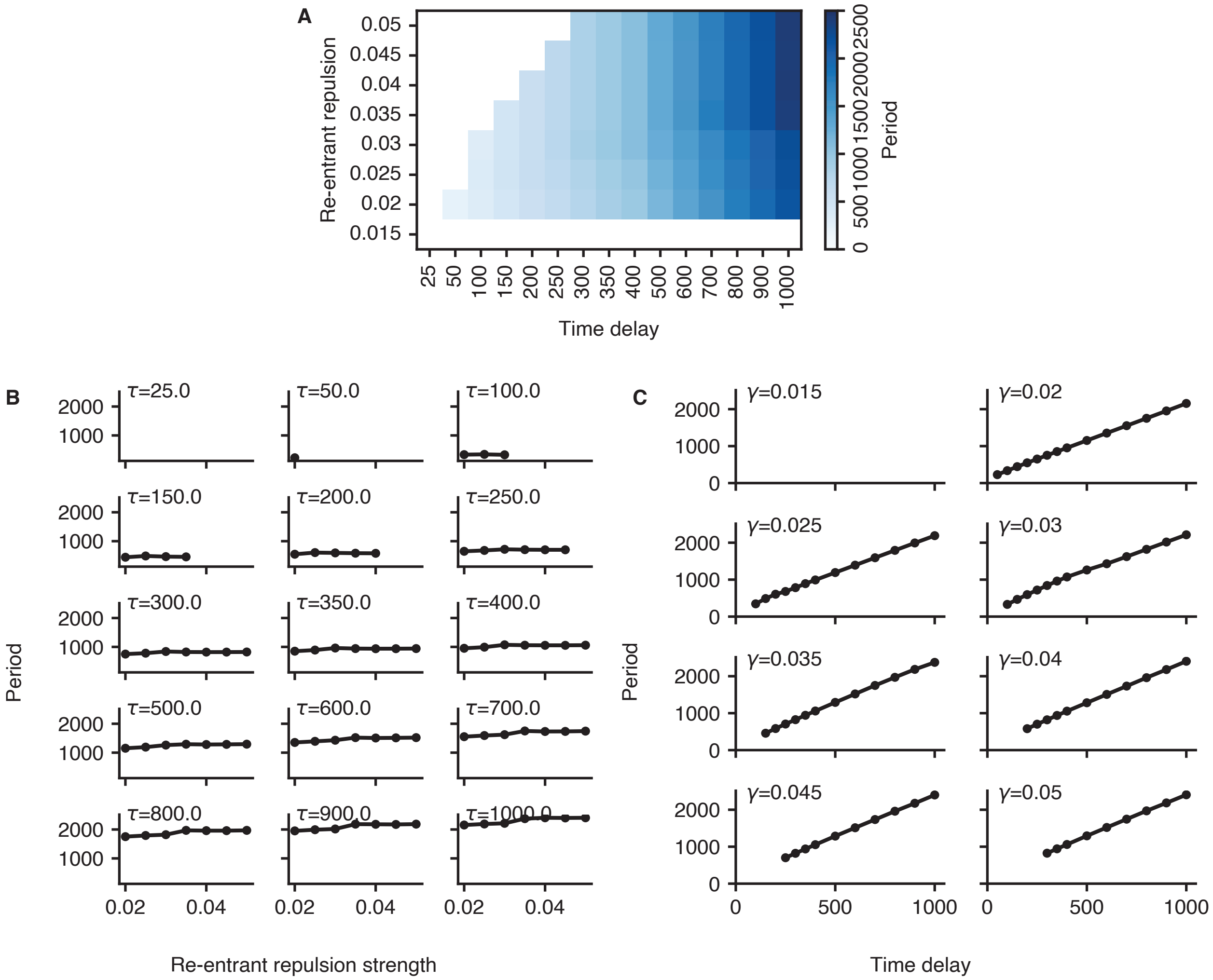


FIG. S12. (A) Period of oscillations depends on the time delay ($\tau$) but is insensitive to the strength of protein-RNA repulsion at high RNA concentrations ($\gamma$). (B) Period of oscillations as a function of the repulsion strength, for fixed time delays $\tau$. (C) Period of oscillations as a function of the time delay, for fixed repulsion strength $\gamma$.

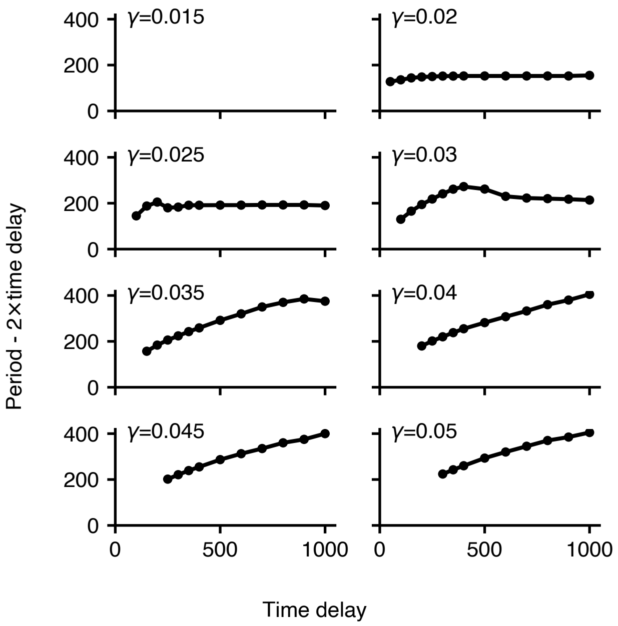


FIG. S13. Period of oscillations minus twice the time delay. This corresponds to the time required to traverse between promoter-proximal and distal positions.

### SV. NON-DIMENSIONALIZATION OF MODEL EQUATIONS

We non-dimensionalize the equations of motion to identify meaningful parameter groupings. Starting from the full dimensional equation,

$$\mathcal{F}(c,m) = \int \mathrm{d}^d r \left[ \frac{\alpha}{4}(c-\bar{c})^4 + \frac{\beta}{2}(c-\bar{c})^2 + \frac{\kappa}{2}|\boldsymbol{\nabla} c|^2 + \frac{\lambda}{2}m^2 + \gamma c m + \frac{\zeta}{2}c^2 m^2 + \eta \exp\left(-\frac{|\boldsymbol{r}-\boldsymbol{r_e}|^2}{2{\sigma_e}^2}\right) c \right], \tag{S29}$$

we scale the concentration fields with the critical concentration $[\mathrm{c}] = \bar{c}$, we scale time with the inverse of the RNA degradation rate $[\mathrm{t}] = 1/k_d$, and scale the spatial dimensions with the RNA diffusion length $[\mathrm{r}] = \sqrt{D_m/k_d}$. We scale the free energy functional using $\alpha[\mathrm{c}]^4[\mathrm{r}]^d$ based on Eq (S29). After scaling the free energy functional, we have

$$\tilde{\mathcal{F}}[\tilde{c},\tilde{m}] = \int \mathrm{d}\tilde{x}\mathrm{d}\tilde{y} \left[ \frac{1}{4}(\tilde{c}-1)^4 + \frac{1}{2}\frac{\beta}{\alpha\bar{c}^2}(\tilde{c}-1)^2 + \frac{1}{2}\frac{\kappa}{\alpha\bar{c}^2(D_m/k_d)}|\tilde{\nabla}\tilde{c}|^2 + \frac{\gamma}{\alpha\bar{c}^2}\tilde{m}\tilde{c} + \frac{1}{2}\frac{\zeta}{\alpha}\tilde{m}^2\tilde{c}^2 + \frac{1}{2}\frac{\lambda}{\alpha\bar{c}^2}\tilde{m}^2 - \frac{\eta}{\alpha\bar{c}^3}\exp\left(\frac{-|\tilde{\boldsymbol{r}}-\tilde{\boldsymbol{r}}_{\boldsymbol{e}}|^2}{2\tilde{\sigma}_e^2}\right) c \right]. \tag{S30}$$

Scaling the equations of motion leads to

$$\partial_{\tilde{t}}\tilde{c} = \tilde{\boldsymbol{\nabla}} \cdot \left( \frac{\alpha\bar{c}^2 M_c}{D_m} \tilde{\boldsymbol{\nabla}} \frac{\delta\tilde{\mathcal{F}}}{\delta\tilde{c}} \right), \tag{S31}$$

$$\partial_{\tilde{t}}\tilde{m} = \tilde{\nabla}^2\tilde{m} + \frac{k_p}{k_d}\exp\left(-\frac{|\tilde{\boldsymbol{r}}-\tilde{\boldsymbol{r}}_p|^2}{2\tilde{\sigma}_p^2}\right) c - \tilde{m}. \tag{S32}$$

We derive the parameter groups from Eqs. (S30), (S31), and (S32):

$$\tilde{\beta} = \frac{\beta}{\alpha\bar{c}^2}, \quad \tilde{\kappa} = \frac{\kappa}{\alpha\bar{c}^2(D_m/k_d)}, \quad \tilde{\gamma} = \frac{\gamma}{\alpha\bar{c}^2}, \quad \tilde{\zeta} = \frac{1}{2}\frac{\zeta}{\alpha}, \quad \tilde{\lambda} = \frac{1}{2}\frac{\lambda}{\alpha\bar{c}^2}, \quad \tilde{\eta} = \frac{\eta}{\alpha\bar{c}^3}, \quad \tilde{M}_c = \frac{\alpha\bar{c}^2 M_c}{D_m}, \quad \tilde{k}_p = \frac{k_p}{k_d}. \tag{S33}$$

We simulate the dimensionful equations with various parameters set to one, such as $\alpha$, $D_m$, and $k_d$. This effectively sets the RNA diffusion length as the length scale and the inverse of the RNA degradation rate as the time scale. We set $k_d$ to not be one only for Fig. 4. We had to decrease $k_d$ in Fig. 4 to obtain center of mass oscillations by increasing the RNA diffusion length. We could also set $\bar{c}$, the critical concentration of the protein, to one. However, we found that doing this reduced the stability of our simulations because it led to negative concentration values. In all simulations, we instead set $\bar{c} = 4$.

### SVI. CODE AVAILABILITY

All simulation and analysis scripts are available on GitHub in the following repositories:

1. Finite volume simulations (FVM) are at https://github.com/gohdavid/CoupledEPCondensates
2. Brownian Dynamics simulations (BD) are at https://github.com/gohdavid/active-polymers
3. Theoretical calculations (Theory) are at https://github.com/gohdavid/CoupledEPCondensatesTheory
4. Finite Element Simulations (FEM) are at https://github.com/gohdavid/CoupledEPCondensatesFEM

The notebooks used to generate the figures in this work are in each of these repositories. Although we do not include simulation data due to storage limitations, we include the simulation scripts used to generate the data.

| **Figure** | **Notebook** |
|---|---|
| Fig. 1B | `FVM/workspace/01_Analysis/Fig_1B.ipynb` and `FVM/workspace/01_Analysis/Fig_1B_I-II-III.ipynb` |
| Fig. 1C | `Theory/Fig_1C_CSV.ipynb` |
| Fig. 2B | `Theory/Fig_2B-4B.ipynb` |
| Fig. 2C | `BD/Fig_2C-S16.ipynb` |
| Fig. 2D | `BD/Fig_2D-S15.ipynb` |
| Fig. 2E | `FVM/workspace/04_Analysis/Fig_2E.ipynb` |
| Fig. 3B | `FVM/workspace/05_Analysis/Fig_3B-3C.ipynb` |
| Fig. 3C | `FVM/workspace/05_Analysis/Fig_3B-3C.ipynb` |
| Fig. 3D | `FVM/workspace/05_Analysis/Fig_3D.ipynb` |
| Fig. 4B | `Theory/Fig_2B-4B.ipynb` |
| Fig. 4C | `FVM/workspace/05_Analysis/Fig_4C-4D.ipynb` |
| Fig. 4D | `FVM/workspace/05_Analysis/Fig_4C-4D.ipynb` |
| Fig. S1 | `FVM/workspace/00_Misc/Fig_S1-S8-S9-S10.ipynb` |
| Fig. S2 | `Theory/Fig_S2-S3-S5-S6.ipynb` |
| Fig. S3 | `Theory/Fig_S2-S3-S5-S6.ipynb` |
| Fig. S5 | `Theory/Fig_S2-S3-S5-S6.ipynb` |
| Fig. S6 | `Theory/Fig_S2-S3-S5-S6.ipynb` |
| Fig. S7 | `FEM/Fig-S7.ipynb` |
| Fig. S8 | `FVM/workspace/00_Misc/Fig_S1-S8-S9-S10.ipynb` |
| Fig. S9 | `FVM/workspace/00_Misc/Fig_S1-S8-S9-S10.ipynb` |
| Fig. S10 | `FVM/workspace/00_Misc/Fig_S1-S8-S9-S10.ipynb` |
| Fig. S11 | `FVM/workspace/05_Analysis/Fig-S11-S12-S13.ipynb` |
| Fig. S12 | `FVM/workspace/05_Analysis/Fig-S11-S12-S13.ipynb` |
| Fig. S13 | `FVM/workspace/05_Analysis/Fig-S11-S12-S13.ipynb` |
| Fig. S14 | `FVM/Fig_S14.ipynb` |
| Fig. S15 | `BD/Fig_2D-S15.ipynb` |
| Fig. S16 | `BD/Fig_2C-S16.ipynb` |

TABLE S1. Notebooks used to generate figures.

### SVII. INCREASED SENSITIVITY OF TRANSCRIPTION TO PROTEINS

Here, we discuss why we implement a Hill function $f_+$, which increases the sensitivity of the RNA production term to the protein field, in Eq. (15). We use $\bar{c} = 4.0$ and $\tilde{\beta} = -0.25$ [Eq. (1)] in our simulations such that there is a small difference between the light and dense phase binodal points ($\tilde{c}_- = 3.5$ and $\tilde{c}_+ = 4.5$). We found this improves simulation stability, as large differences in protein concentration between dense and light phases can numerically result in negative concentration values. However, when using these parameters in Sec. V, we find that the difference in repulsive forces when the droplet is far and close to the promoter is insufficient to lead to oscillations: to get oscillations, we need repulsions to be weak when the condensate is far, and strong when the condensate is close. This difference in repulsive forces depends on the difference in dense and light phase concentrations: when the condensate is at the promoter, it increases RNA production proportional to the change in protein concentration ($\tilde{c}_+ - \tilde{c}_-$) [Eqs. (5) and (15)], resulting in increased RNA concentrations and subsequently increased repulsions [Eq. (2)]. Because $\tilde{c}_+$ and $\tilde{c}_-$ are similar in our simulations for numerical reasons, we need to use $f_+$ to increase the change in RNA production and exhibit oscillations. We implement a Hill function centered at the critical point $\bar{c} = 4$ by using the parameters $\bar{c} = 3$, $K_d = 1$, and $n = 2$:

$$f_+(c) = \frac{V_{\max}(c - \bar{c})^n}{(c - \bar{c})^n + K_d^n} \,. \tag{S34}$$

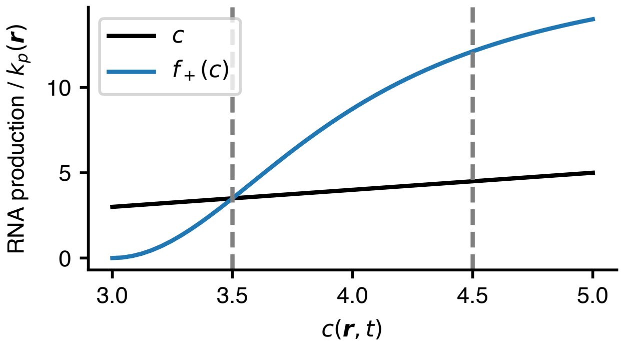


FIG. S14. We increase the sensitivity of the production rate to the concentration of the protein using a Hill function $f_+$.

The parameter $V_{\max}$ can be subsumed into $k_p$, but we explicitly specify $V_{\max} = 17.5$, which sets the production rate in the light phase to be the same with and without $f_+$. This makes it easier to compare $k_p$ values (Fig. S14).

### SVIII. UNITS AND CONVERSIONS

In the present section, we outline the estimates of parameters that we use in our Brownian dynamics simulations of a Rouse chain. We convert the length of chromatin from bp to nm using the following relation, which assumes an average linker length of bare DNA connecting adjacent nuclesosomes of 45 bp [5],

$$L_c[\text{bp}] = L_c[\text{nm}] \times \frac{0.34\,\text{bp}}{\text{nm}} \times \frac{45 \text{ bp (linker)}}{45 + 146 \text{ bp (linker + nucleosome)}}. \tag{S35}$$

For example, we estimate that an enhancer and promoter separated by 100 kbp would have a contour length of 2000 nm. We choose a Kuhn length based on predictions from a "zig-zag" polymer model of nucleosome-bound DNA in mouse embryonic stem cells [5],

$$b = 35.36\,\text{nm} \approx 441.42\,\text{bp}. \tag{S36}$$

Thus, for a given enhancer-promoter genomic separation, we can calculate the number of monomers $N$ in our polymer chain to be $L_c/b$. The associated Rouse time for that polymer would then be $\tau_R = L_c^2/(3\pi^2 D_{\text{chrom}})$. We calculate the diffusion coefficient of chromatin from its apparent diffusion coefficient $D_{\text{app}} \approx 0.01\,\mu\text{m}^2\,\text{s}^{-1/2}$ [6],

$$D_{\text{chrom}} = \left[\frac{D_{\text{app}}}{(12/\pi)^{1/2} b}\right]^2 = \frac{\pi D_{\text{app}}^2}{12 b^2} \approx 0.02\,\mu\text{m}^2\,\text{s}^{-1}. \tag{S37}$$

For the diffusion coefficient of nuclear RNA [7], we take

$$D_{\text{RNA}} \approx 0.03\,\mu\text{m}^2\,\text{s}^{-1} \text{ to } 0.1\,\mu\text{m}^2\,\text{s}^{-1}\,. \tag{S38}$$

For the average radius of a condensate [8, 9], we take

$$R_{\text{droplet}} = 250\,\text{nm}\,. \tag{S39}$$

For the lifetime of nascent RNA [10], we take

$$k_d^{-1} \approx 10\,\text{min} \tag{S40}$$

For the diffusion length of RNA, we take

$$\ell = \sqrt{\frac{D_{\text{RNA}}}{k_d}} = \sqrt{\frac{0.03\,\mu\text{m}^2}{\text{s}} \times 10\,\text{min} \times \frac{60\,\text{s}}{1\,\text{min}}} = 4.24\,\mu\text{m}\,. \tag{S41}$$

In our polymer simulations, we set $b = 1$ and $D_{\text{chrom}} = 1$. Hence, the Kuhn length and the diffusion coefficient of chromatin set the length and time scales of our simulations,

$$L \sim b, \tag{S42}$$

$$t \sim b^2/D_{\text{chrom}}. \tag{S43}$$

In Fig. 2B, we calculated the dimensionless velocity $\tilde{v}$ using unitful values of $R_{\text{droplet}}$ and $\ell$. In Fig. 2C, we convert the distance to experimental units by scaling it with $L$. In Fig. 2D, we convert the velocity scale $\nu$ to experimental units by scaling it with $L/t$,

$$L/t \sim D_{\text{chrom}}/b \approx 0.592\,\mu\text{m}\,\text{s}^{-1}, \tag{S44}$$

and convert the linear genomic distance to experimental units by scaling with $L$.

### SIX. ENHANCER-PROMOTER CONTACT PROBABILITY IS MORE SENSITIVE TO CHANGES IN CONDENSATE VELOCITY FOR CLOSE PAIRS

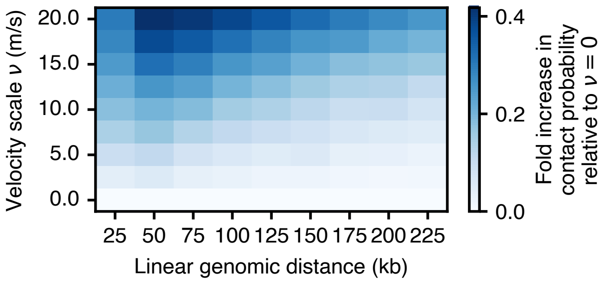


FIG. S15. The fold increase in enhancer-promoter contact probability relative to $\nu = 0$, as a function of linear distance between the two loci and the scale of the condensate velocity towards the promoter. Contact radius is 250 nm. Enhancer-promoter contact probability is more sensitive to changes in condensate velocity for close enhancer-promoter pairs as compared to far enhancer-promoter pairs.

### SX. EFFECT OF CHANGING ACTIVITY AND FRICTION OF THE ENHANCER

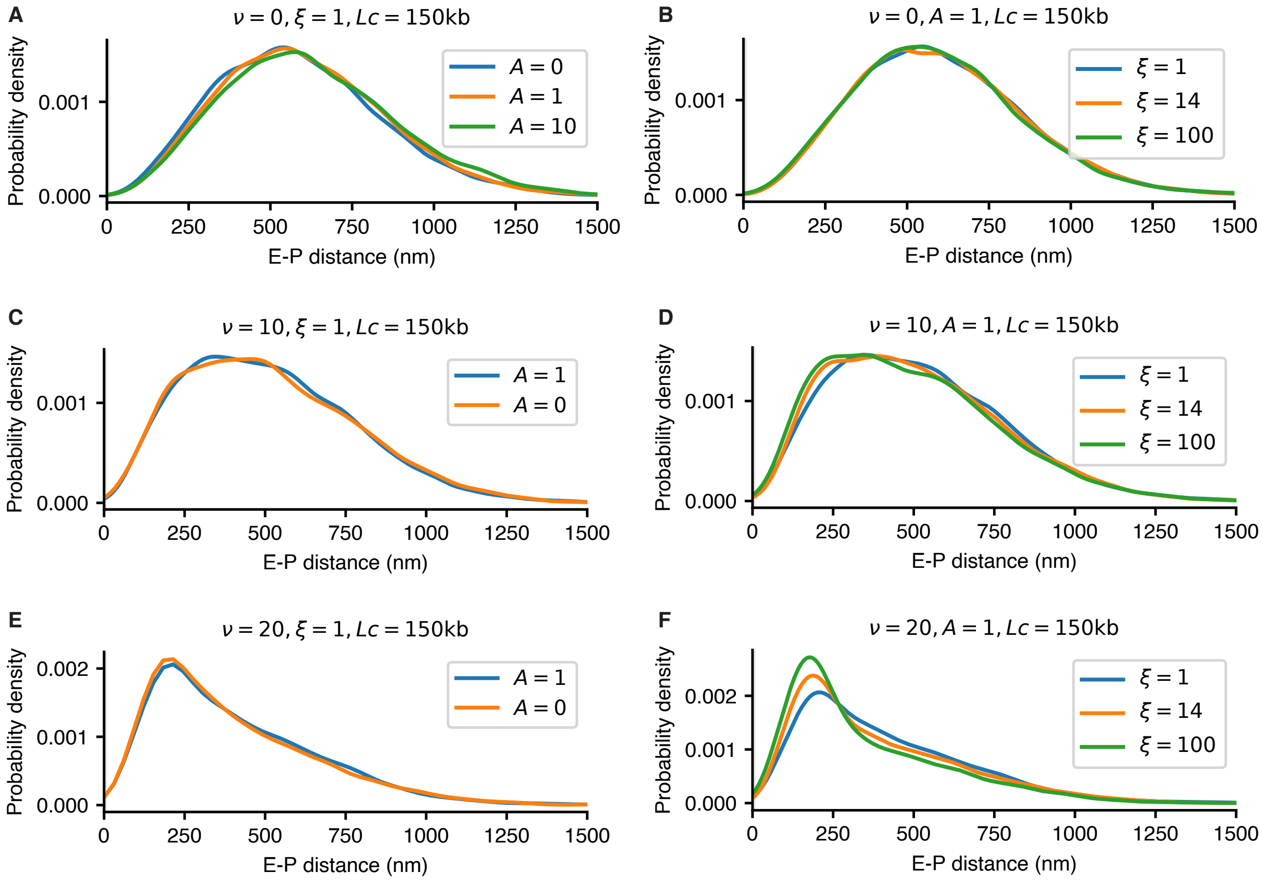


FIG. S16. Enhancer-promoter distance distribution calculated using various values of the velocity scale $\nu$ for different values of the activity $A$ and friction $\xi$ of the enhancer monomer. Only for large values of $\nu$ [panel (F)], which corresponds to strong non-reciprocal interactions between enhancer and promoter, do we observe that the friction of the enhancer has an effect on the steady-state.

---


[1] C. R. Harris, K. J. Millman, S. J. van der Walt, R. Gommers, P. Virtanen, D. Cournapeau, E. Wieser, J. Taylor, S. Berg, N. J. Smith, R. Kern, M. Picus, S. Hoyer, M. H. van Kerkwijk, M. Brett, A. Haldane, J. F. del Río, M. Wiebe, P. Peterson, P. Gérard-Marchant, K. Sheppard, T. Reddy, W. Weckesser, H. Abbasi, C. Gohlke, and T. E. Oliphant, Nature **585**, 357 (2020).

[2] P. Virtanen, R. Gommers, T. E. Oliphant, M. Haberland, T. Reddy, D. Cournapeau, E. Burovski, P. Peterson, W. Weckesser, J. Bright, S. J. van der Walt, M. Brett, J. Wilson, K. J. Millman, N. Mayorov, A. R. J. Nelson, E. Jones, R. Kern, E. Larson, C. J. Carey, İ. Polat, Y. Feng, E. W. Moore, J. VanderPlas, D. Laxalde, J. Perktold, R. Cimrman, I. Henriksen, E. A. Quintero, C. R. Harris, A. M. Archibald, A. H. Ribeiro, F. Pedregosa, and P. van Mulbregt, Nature Methods **17**, 261 (2020).

[3] A. Bray, Advances in Physics **43**, 357 (1994).

[4] A. Goychuk, L. Demarchi, I. Maryshev, and E. Frey, Physical Review Research **6**, 033082 (2024).

[5] B. Beltran, D. Kannan, Q. MacPherson, and A. J. Spakowitz, Physical Review Letters **123**, 208103 (2019).

[6] B. Gu, T. Swigut, A. Spencley, M. R. Bauer, M. Chung, T. Meyer, and J. Wysocka, Science (New York, N.Y.) **359**, 1050 (2018).

[7] T. Misteli, Histochemistry and Cell Biology **129**, 5 (2008).

[8] W.-K. Cho, J.-H. Spille, M. Hecht, C. Lee, C. Li, V. Grube, and I. I. Cisse, Science **361**, 412 (2018).

[9] M. Du, S. H. Stitzinger, J.-H. Spille, W.-K. Cho, C. Lee, M. Hijaz, A. Quintana, and I. I. Cissé, Cell **187**, 331 (2024).
[10] T. Muramoto, D. Cannon, M. Gierliński, A. Corrigan, G. J. Barton, and J. R. Chubb, Proceedings of the National Academy of Sciences **109**, 7350 (2012).
